# Dimerization of ADAR1 modulates site-specificity of RNA editing

**DOI:** 10.1101/2023.12.05.570066

**Authors:** Allegra Mboukou, Vinod Rajendra, Serafina Messmer, Marjorie Catala, Carine Tisné, Michael F. Jantsch, Pierre Barraud

**Author notes:** AM and VR contributed equally to this work. Corresponding authors: Michael F. Jantsch, or Pierre Barraud.

## Abstract

Adenosine-to-inosine editing is catalyzed by adenosine deaminases acting on RNA (ADARs) in double-stranded RNA (dsRNA) regions. Although three ADARs exist in mammals, ADAR1 is responsible for the vast majority of the editing events and acts on thousands of sites in the human transcriptome. ADAR1 has been proposed to form a stable homodimer and dimerization is suggested to be important for editing activity. In the absence of a structural basis for the dimerization of ADAR1, and without a way to prevent dimer formation, the effect of dimerization on enzyme activity or site specificity has remained elusive. Here, we report on the structural analysis of the third double-stranded RNA-binding domain of ADAR1 (dsRBD3), which reveals stable dimer formation through a large inter-domain interface. Exploiting these structural insights, we engineered an interface-mutant disrupting ADAR1-dsRBD3 dimerization. Notably, dimerization disruption did not abrogate ADAR1 editing activity but intricately affected editing efficiency at selected sites. This suggests a complex role for dimerization in the selection of editing sites by ADARs, and makes dimerization a potential target for modulating ADAR1 editing activity.

## INTRODUCTION

Adenosine deaminases acting on RNA (ADARs) convert adenosines (A) to inosines (I) in double-stranded endogenous and viral RNAs. Inosines are interpreted as guanosines by most cellular machineries and therefore A-I conversions can recode mRNAs but can also affect the folding or antigenicity of RNAs. Two genes that produce active ADAR proteins can be found in mammals, *Adar* and *Adarb1* [1]. *Adarb1* gives rise to the ADAR2 protein which localizes to nuclei and is mainly expressed in the nervous system, the cardiovascular system and the gastrointestinal tract [2,3]. In mice, loss of ADAR2 leads to lethality around day 21 which can be rescued by pre-editing one of its key substrates, the glutamate receptor subunit *Gria2* [4]. *Adar* encodes the ADAR1 protein which is expressed in all tissues and comes in two versions. A 150kDa protein (ADAR1 p150) is expressed from an interferon inducible promoter and is mainly localized to the cytoplasm. A constitutively expressed ADAR1 p110 version lacks the amino terminus of p150 and is mainly localized to the nucleus [5,6]. Both ADAR1 p110 and p150 share a C-terminal deaminase domain and three double-stranded RNA-binding domains (dsRBDs) required for substrate binding. A nuclear localization signal made of two modules is flanking the third dsRBD in both ADAR1 isoforms [7]. The amino terminus of ADAR1 p150 harbors two Z-DNA binding domains and a nuclear export signal [8,9]. Interestingly, both ADAR1 p150 and p110 can shuttle between the nucleus and the cytoplasm although the steady state localization of p150 is cytoplasmic and that of p110 is nuclear [10,11].

Loss of ADAR1 p150 leads to embryonic lethality while loss of ADAR1 p110 leads to post-natal runted appearance [12]. Interestingly, specific mutations within ADAR1 p150 suggest that different domains contribute to different pathways and phenotypes [13]. Loss of RNA editing activates the innate immune sensor MDA5 leading to interferon signaling [14–16]. The dsRBDs seemingly compete for dsRNA binding with PKR and prevent its activation [17,18]. The ZBDs of ADAR1 p150 have been shown to prevent the activation of ZBP1, another Z-RNA binding protein that leads to necroptosis upon Z-RNA binding [19–21].

Upon binding to double-stranded RNA substrates, the catalytic domain of ADARs flips the substrate adenosine out of its double-stranded context to allow the hydrolytic deamination to occur [22]. As base-flipping is inhibited by base-pairing, ADARs act most efficiently on unpaired adenosines within a certain sequence context [23]. Still, neither dsRBDs nor the catalytic domain show a strong sequence specificity and consequently extended stretches of dsRNA can be edited rather promiscuously. Interestingly, however, RNA-editing events that lead to protein recoding can occur very site-specifically. Structural features of dsRNA substrates seemingly help to correctly position the dsRBDs and the catalytic domain of ADARs. Additionally, dimer formation of ADARs has repeatedly been observed and was discussed as a way to facilitate correct substrate recognition [24–26]. A crystal structure of ADAR2 dsRBD2 and deaminase domains in complex with RNA revealed that this protein can form an asymmetric dimer via its catalytic deaminase domain [27]. Similarly, *Drosophila* ADAR can form dimers albeit via its N-terminal region [28]. Dimer formation of mammalian ADAR1 proteins has also been demonstrated by co-IP and size fractionation experiments [29], as well as fluorescence resonance energy transfer (FRET) experiments in cells transfected with fluorophore-bearing ADAR1 [30]. While this dimer formation was shown to be independent of the protein’s RNA binding ability, the precise region for ADAR1 homodimerization remained enigmatic [31].

Double-stranded RNA-binding domains have been shown to allow protein-protein interactions and dimer formation [32]. In ADAR1, a region containing its third dsRBD (dsRBD3) had been shown to be essential for dimer formation [33]. In order to identify the exact elements involved in the dimerization of ADAR1, and to characterize the underlying structural aspects, we set out to understand this domain in more detail using X-ray crystallography and solution scattering. Here, we report on the structural analysis of ADAR1-dsRBD3, which reveals dimer formation with a large interdomain interface at the level of the dsRBD’s β-sheets. In addition, co-immunoprecipitation and mutational analysis confirm a dsRBD3-mediated dimer formation of ADAR1 *in vivo*. We show further, that dimer formation affects editing efficiency at selected sites in ADAR1 model substrates, suggesting a complex role for dimerization in the selection of editing sites by ADARs.

## RESULTS

### The folded region of ADAR1-dsRBD3 is sufficient to mediate dimerization

Nishikura and colleagues identified that regions around dsRBD3 of ADAR1 are required for the homodimerization of ADAR1 [33]. More precisely, a construct lacking the segment covering residues 725-833 resulted in the loss of interaction with full-length ADAR1. We determined earlier that the folded part of dsRBD3 of ADAR1 consists of residues 716-797 [7]. We therefore wanted first to better delineate the elements needed for ADAR1 dimerization and to determine whether the folded domain or the flanking regions, or both, are involved in the interaction. To do so, we produced and purified three constructs of different size, which all encompass at least the folded part of ADAR1-dsRBD3 (*i.e.* residues 716-797), but with different flanking regions, namely dsRBD3-long (residues 688-817), dsRBD3-mid (residues 708-801) and dsRBD3-short (residues 716-797) (Supplementary Figure S1A). Purification tags were cleaved off to avoid potential unspecific interactions. All constructs were analyzed by small angle X-ray scattering experiments coupled to size exclusion chromatography (SEC-SAXS). The scattering data revealed that all three constructs behave as dimeric domains in solution (Table 1 and Supplementary Figure S1B). These experiments demonstrate that ADAR1 dimerization is an intrinsic property of its third dsRBD that forms a stable dimer in solution on its own, without the need for any additional domain.

Table 1.

**Table 1**
| ADAR1-dsRBD3 construct | MW (kDa) | R<sub>g</sub> (Å) | D<sub>max</sub> (Å) | apparent MW [95%]<sup>a</sup> (kDa) |
| --- | --- | --- | --- | --- |
| dsRBD3-long | 14.5 | 25.5 ± 1.0 | 80 | 30.9 [29.2 - 33.4] |
| dsRBD3-mid | 10.8 | 19.8 ± 1.2 | 66 | 19.9 [19.0 - 20.9] |
| dsRBD3-short | 9.2 | 19.2 ± 1.1 | 55 | 21.2 [19.6 - 22.7] |

<sup>a</sup> the apparent MW in solution is given as an averaged value as well as an interval of confidence at 95% in brackets, all determined with PRIMUS (ATSAS suite) using default settings.

### The crystal structure of ADAR1-dsRBD3 reveals a dimerization interface

We determined the crystal structure of the folded entity of ADAR1-dsRBD3 (residues 716-797) and refined it to a resolution of 1.65 Å (Supplementary Table S1). ADAR1-dsRBD3 crystallized in the hexagonal space group P3_1_21 with two dsRBDs in the asymmetric unit (Figure 1A). Overall, the structure of the individual dsRBDs corroborate the structure previously solved by NMR spectroscopy, confirming the classical topology of dsRBDs with the core α1-β1-β2-β3-α2 elements, as well as the additional N-terminal helix α_N_, important for assembling the bimodular NLS of ADAR1 [7]. However, as a consequence of the way NMR structures are calculated, namely by collecting and interpreting a large number of short-range distances, the previously solved solution structure failed at identifying the dimeric state of the domain. This most probably also arose since ADAR1-dsRBD3 forms a symmetric dimer, in which the NMR resonances coming from identical residues in each monomer are perfectly superimposable. On the contrary, in a crystal structure, the overall assembly is readily accessible, which here revealed that the two dsRBDs of the asymmetric unit interact via an extended interface of ∼490 Å^2^ at the level their β-sheet surface (Figure 1). The two monomer structures are not perfectly identical as a result of small local variations caused by differences in the packing assembly, and display an r.m.s.d. of 1.19 Å over the entire Cα of the domains (Supplementary Figure S2). Nevertheless, the monomers are assembled via a pseudo C2-symmetry and form a virtually symmetric dimer (Figure 1B), that would most probably be truly symmetric without the structural distortions caused by crystal packing. Interactions at the dimer interface involve residues from all β-strands, namely V747 and D748 from β1, K757, V759 and Q761 from β2, and W768, P770, A771 and C773 from β3. Interactions are formed by a combination of electrostatic and hydrophobic contacts. On one edge of the β-sheet, W768 stacks onto the P770 cycle and the side chain Q761 interacts via two hydrogen bonds with the backbone of β3, namely the carbonyl of F769 and the amide of A771 (Figure 1C). On the other edge, a small hydrophobic core is assembled via the side chains of V747, V759 and C773, and an electrostatic contact is formed between D748 and K757 (Figure 1D). All contacts are present twice between each of the monomers owing to the symmetrical nature of the interface.

**Figure 1:**
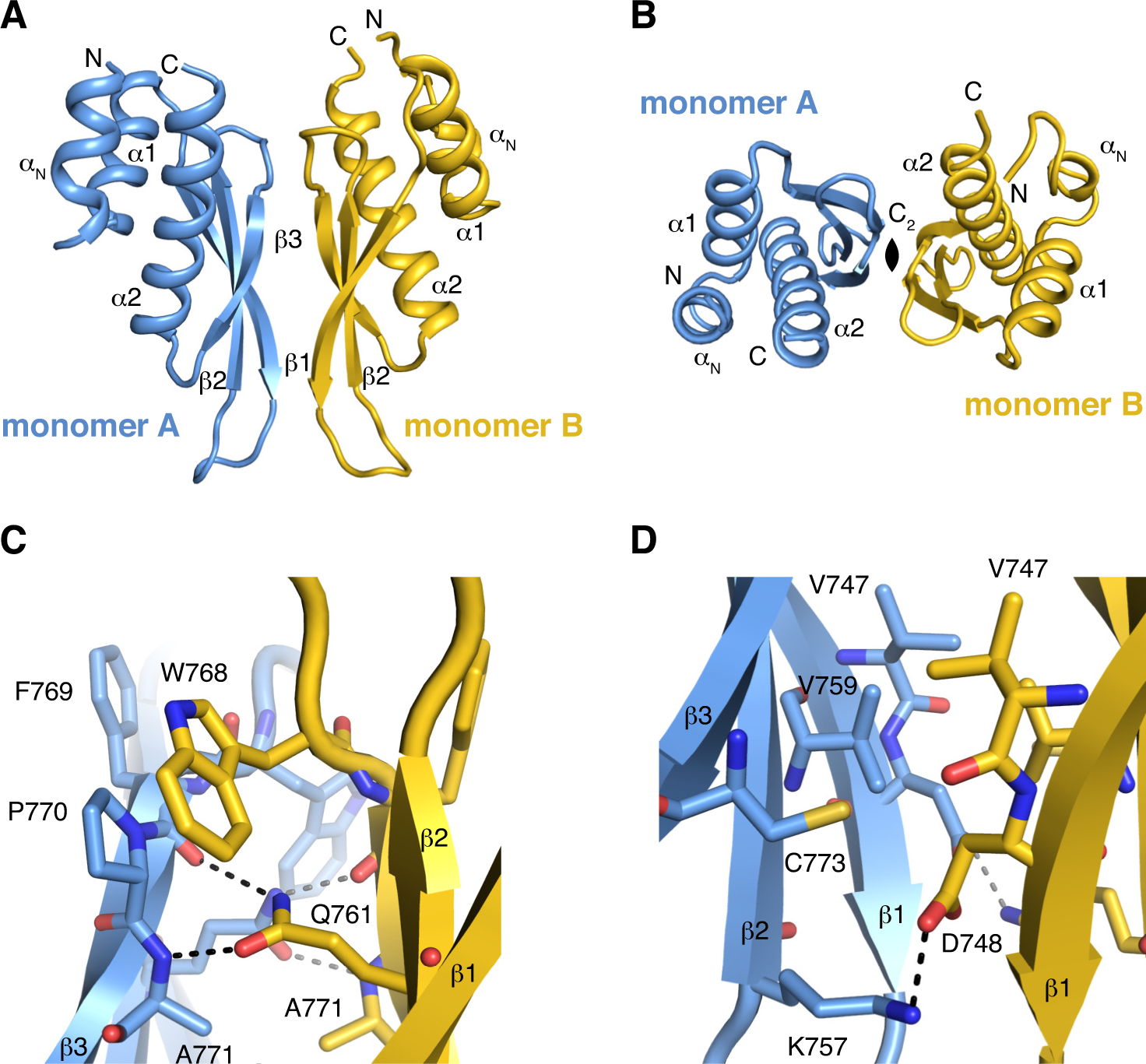
ADAR1-dsRBD3 forms a stable symmetric dimer. (**A**) Overall organization of ADAR1-dsRBD3 dimer. Monomers A and B are displayed in cartoon mode and coloured *in blue* and *in yellow*, respectively. Secondary structure elements are labelled on each monomer. (**B**) ADAR1-dsRBD3 forms a symmetric dimer. Top view along the C_2_ symmetry axe. (**C**) Contacts at the dimer interface between monomer A (*in blue*) and monomer B (*in yellow*) on one edge of the β-sheet. Important residues are shown as sticks. Polar contacts are shown as dashed lines. (**D**) Contacts at the dimer interface between monomer A (*in blue*) and monomer B (*in yellow*) on the other edge of the β-sheet. Important residues are shown as sticks. Polar contacts are shown as dashed lines.

### The dimer observed in the crystal corresponds to the one existing in solution

Next, to establish that the dimer observed in the crystal structure is also the one that exist in solution, we calculated theoretical scattering curves from the crystal structure of ADAR1-dsRBD3 using CRYSOL and compared them with the experimental SAXS data obtained with the dsRBD3-short construct. The theoretical curve obtained with the ADAR1-dsRBD3 monomer structure differs substantially from the experimental data (goodness of fit χ^2^ = 41.8), but the one obtained with the ADAR1-dsRBD3 dimer is highly similar to the experimental data (goodness of fit χ^2^ = 1.30) (Figure 2). This indicates that the dimeric structure observed in the crystal is also a good representation of ADAR1-dsRBD3 in solution.

**Figure 2:**
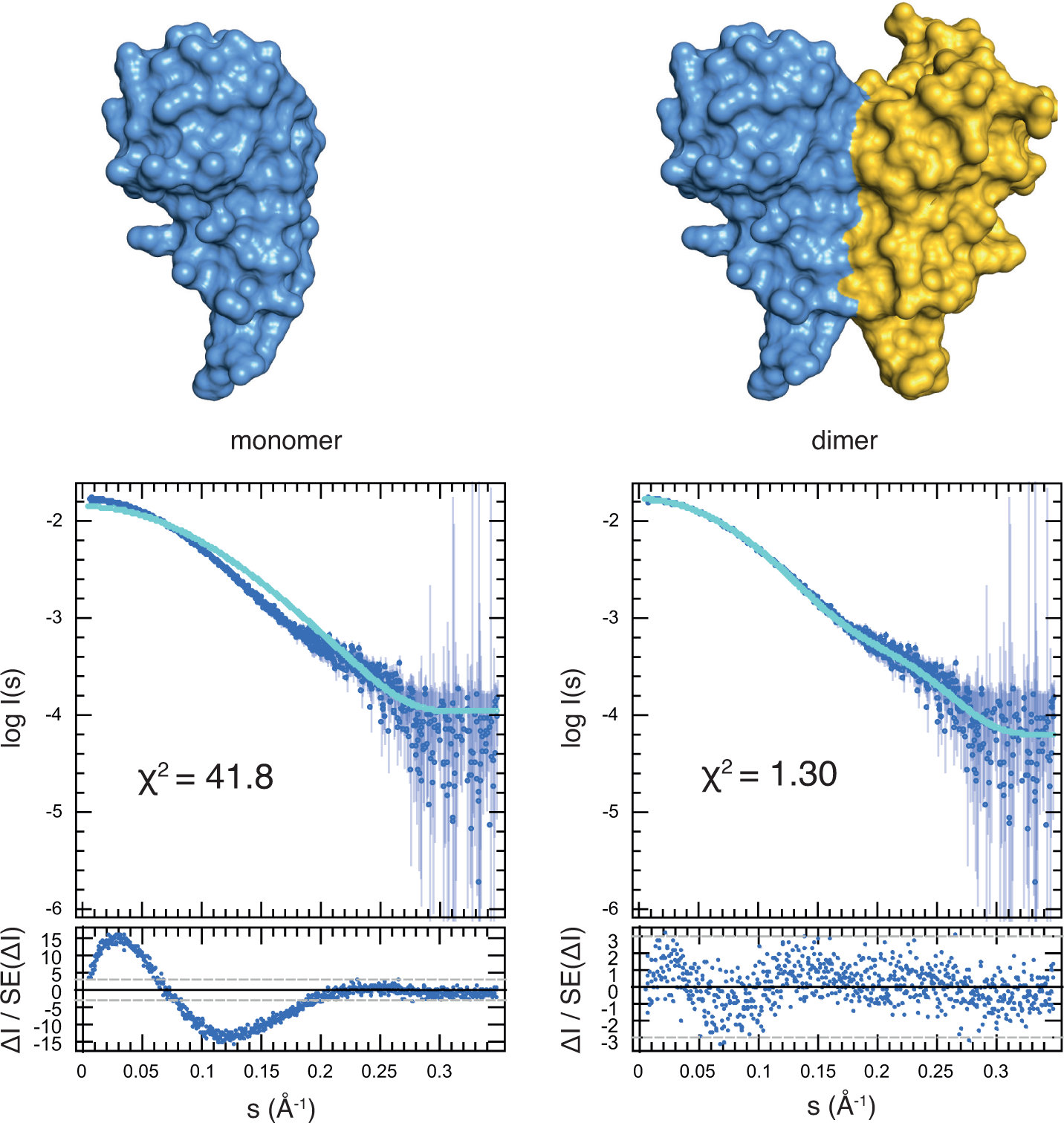
The ADAR1-dsRBD3 dimer observed in the crystal also exists in solution. SAXS characterization of ADAR1-dsRBD3 (dsRBD3-short construct, residues 716-797). Upper panel: Surface representation of ADAR1-dsRBD3 monomer (*left*) or dimer (*right*) structure determined by crystallography (see Figure 1). Lower panel: Experimental and theoretical scattering curve calculated from the crystal structure atomic coordinates of dimeric and monomeric ADAR1-dsRBD3, along with their residuals, as reported by CRYSOL. Goodness of fit (χ^2^) are reported in each case.

### Dimerization of ADAR1-dsRBD3 is compatible with its binding to dsRNA

Having shown that ADAR1-dsRBD3 dimerizes via its entire β-sheet surface, we were wondering whether this dimerization affects the binding of the domain to dsRNA. We were indeed wondering whether the strong binding of ADAR1-dsRBD3 to dsRNA that we previously characterized by ITC [7] occurs with ADAR1-dsRBD3 in its dimeric form. For that, we sought at obtaining structural information to reveal how ADAR1-dsRBD3 interacts with dsRNA. For this purpose, we have set up crystallization assays using small oligonucleotides of 11-13 residues that can base-pair with themselves to form short duplexes and associate with other duplexes via their two-nucleotide overhangs to form long dsRNA helices, a strategy designed to facilitate crystallization [34]. We could obtain well-diffracting crystals only with the longest RNA, *e.g.* 13-nucleotide long RNA, and refined the structure of the ADAR1-dsRBD3/dsRNA complex to a resolution of 2.8 Å (Supplementary Table S1). ADAR1-dsRBD3 crystallized in the hexagonal space group P6_1_22 with two dsRBDs and three oligonucleotides in the asymmetric unit (Figure 3A). Within the crystal, the oligonucleotides are assembled into long pseudo-A-form RNA helices through their CG 3’-overhangs (Figure 3B). Each ADAR1-dsRBD3 monomer binds to such a dsRNA helix at the exact same position, meaning that residues from each dsRBD are forming the same type of contacts with identical nucleotides on two different dsRNA helices. These contacts include mainly non-sequence-specific contacts with the sugar-phosphate backbone of the RNA helices (Figure 3C). Typically, molecular recognition of dsRNA by dsRBDs occurs via three regions of interaction: helix α1 and the loop between β1 and β2 contact two dsRNA minor grooves at one turn of interval, whereas the N-terminal tip of helix α2 contacts the dsRNA phosphate backbone across the major groove [35]. The binding mode that we observe here is similar to what was observed for other dsRBD-dsRNA complexes [36–44]. In particular, in region 1, helix α1 makes non-sequence-specific contacts with 2’-OH groups on each RNA strands, namely with U9 and A4, via residues E733 and R736, respectively (Figure 3C, D). A point worth noting in this region concerns the additional N-terminal helix α_N_ that also participates in the recognition by making additional contacts with the sugar-phosphate backbone, namely with U8 and G5, via residues R721 and N726, respectively (Figure 3C, D). In region 2, residue H754 forms non-sequence-specific contacts with 2’-OH groups on each RNA strands, namely with C10 and A3, via its main-chain carbonyl and its imidazole ring Nδ1, respectively. In addition, residue P753 forms a sequence-specific contact with the exocyclic amino group of G2, via its main-chain carbonyl (Figure 3C, E), as previously reported in many dsRBD-dsRNA structures [35]. Finally, in region 3, lysines K777 and K778 form electrostatic contacts with the phosphate backbone of each RNA strand across the major groove. Namely, K777 interacts with non-bridging oxygen atoms of C12 and G13, via its main-chain amide and side-chain terminal amino group, respectively; and K778 interacts with non-bridging oxygen atoms of C7 and U8 on the other strand, via its side-chain terminal amino group. Additionally, residue Q782 contacts the 3’ bridging oxygen atom of C6 via its side-chain amide group (Figure 3C, F). Finally, the third lysine of the conserved KKxxK motif, namely K781, is found further away from the RNA sugar-phosphate backbone (> 6 Å), but could form a water-mediated contact with a non-bridging oxygen atom of C6, as previously reported [45]. Altogether, apart from the remarkable contribution of helix α_N_, RNA-binding by individual monomers of ADAR1-dsRBD3 corresponds to the well-defined canonical mode of dsRNA binding.

**Figure 3:**
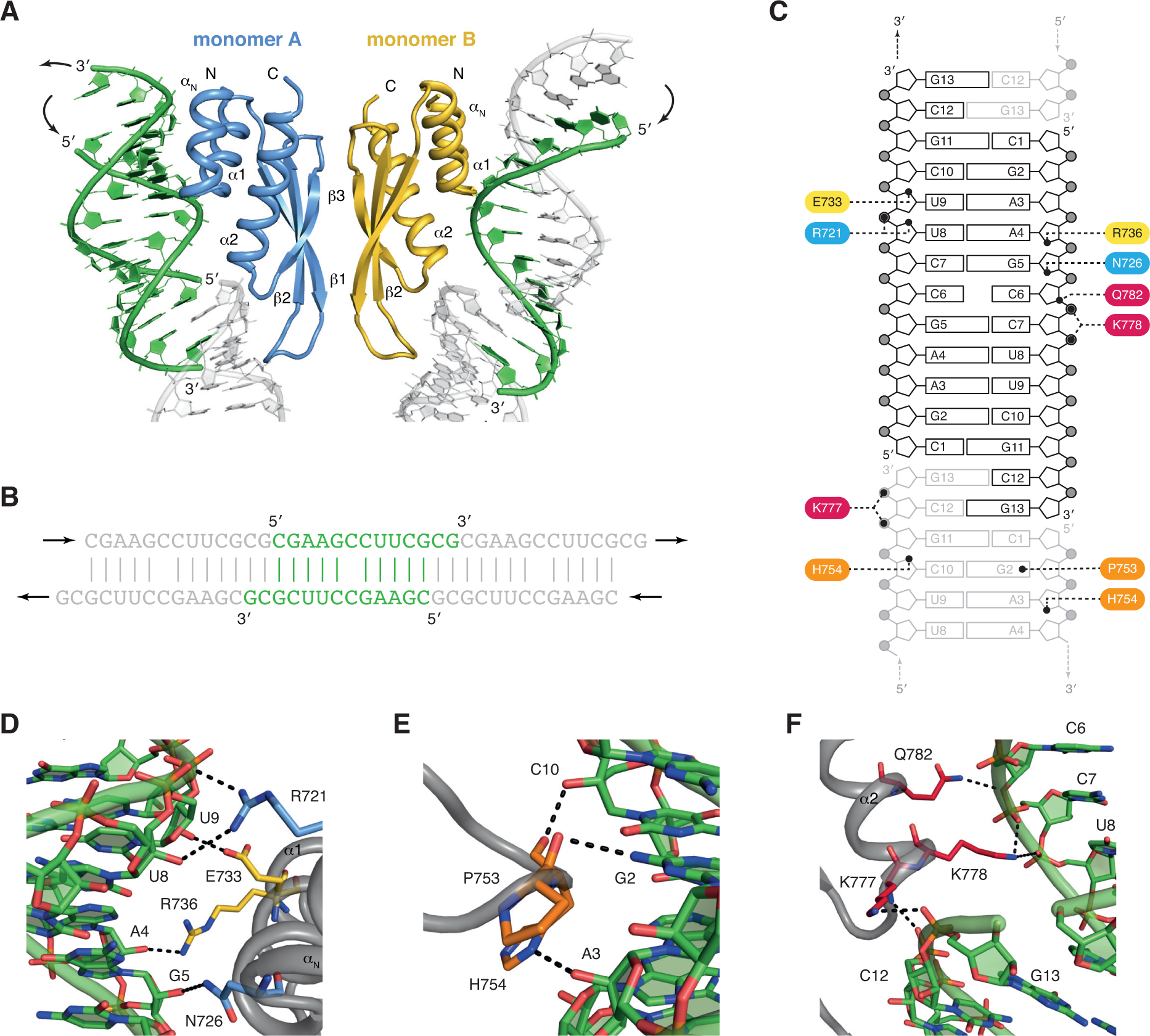
Dimerization of ADAR1-dsRBD3 is compatible with its binding to dsRNA. (**A**) Overall organization of the asymmetric unit of ADAR1-dsRBD3:dsRNA crystal structure. Monomers A and B are displayed in cartoon mode and coloured *in blue* and *in yellow*, respectively. Secondary structure elements are labelled on each monomer. dsRNA helices are displayed in green and in grey, for RNA strands within the asymmetric unit or symmetry-related RNA strands, respectively. (**B**) Schematic representation of the RNA self-assembly. The RNA sequence (5’-CGAAGCCUUCGCG-3’) contains two-nucleotides 3’-overhangs used to generate a dsRNA by self-hybridization. The central dsRNA block is shown *in green*, and the flanking blocks *in grey*. Arrows indicate the direction of self-assembly that leads to a pseudo-A-form dsRNA helix. (**C**) Schematic representation of ADAR1-dsRBD3:dsRNA contacts. Dotted lines indicate contacts between ADAR1-dsRBD3 residues and the RNA. Residues from various dsRBD regions are shown with different colours, with residues from helix αN *in blue*, from helix α1 *in yellow*, from the β1-β2 loop *in orange*, and from the N-terminal tip of helix α2 *in red*. (**D**–**F**) Detailed views of the three regions of interactions. Polar contacts are displayed as dotted lines. The same colour code is used as in panel C.

In addition to the RNA recognition mode by individual monomers, in the case of ADAR1-dsRBD3, the ability to bind to two dsRNA helices as a dimer is of particular interest (Figure 3A). Importantly, the two dimers observed in the crystal structures of ADAR1-dsRBD3 alone (Figure 1A), or in complex with dsRNA (Figure 3A), are completely superimposable (r.m.s.d. of 0.63 Å over the entire Cα of the domains – Supplementary Figure S3). Since the dimerization of ADAR1-dsRBD3 occurs via the β-sheet surface, which is located opposite to the dsRNA-binding surface, the dimerization does not prevent the domain to bind to dsRNA, and each dsRBD monomer can bind to a dsRNA helix (Figure 3A). This feature gives ADAR1 an outstanding position in terms of substrate RNA-binding (see discussion).

### Mutations at the interface render ADAR1-dsRBD3 monomeric

In order to investigate the importance of the ADAR1 dimeric status for its cellular functions, we sought to design point mutations at the interface of ADAR1-dsRBD3s that would disrupt the domain interactions and result in a monomeric ADAR1. For this, we have based our strategy on the structural comparison of ADAR1-dsRBD3 with an archetypal monomeric dsRBD, i.e. the Xlrbpa-dsRBD2 (pdb code 1di2) [36], a dsRBD we already used in our previous study to design a chimeric ADAR1-dsRBD3/Xlrbpa-dsRBD2 that retained the nuclear localization properties of ADAR1 [7]. We thus first superimposed our ADAR1-dsRBD3 structure with that of Xlrbpa-dsRBD2 and derived the corresponding structure-based sequence alignment (Supplementary Figure S4). Based on these structure and sequence alignments, and considering the main interactions observed at the ADAR1-dsRBD3 interface (Figure 1C, D), we decided to create a mutated construct with four point-mutations, i.e. V747A and D748Q in strand β1 and W768V and C773S in strand β3, with the aim of disrupting the dimer interface. We expressed and purified this construct and measured its apparent molecular weight with SEC-MALLS and SAXS. As a control, we also studied the chimeric ADAR1-dsRBD3/Xlrbpa-dsRBD2 construct we previously studied [7]. This construct consists of the entire Xlrbpa-dsRBD2 sequence, flanked by the N- and C-terminal modules forming the bimodular NLS of ADAR1, together with the additional N-terminal helix α_N_ of ADAR1 and few point mutations to allow the correct accommodation of this helix (Supplementary Figure S4) [7]. This construct was chosen since it is expected to be monomeric, similar to Xlrbpa-dsRBD2, while retaining the intracellular localization properties of ADAR1. All constructs used in this part were based on dsRBD3-mid (residues 708-801) that retained the N-terminal purification tag, and were checked to be properly folded using NMR spectroscopy (Supplementary Figure S5). Scattering data in solution, i.e. SEC-MALLS and SAXS, confirm that the chimeric ADAR1-dsRBD3/Xlrbpa-dsRBD2 construct is indeed monomeric (Table 2 and Supplementary Figure S6). Most importantly, the mutated construct at the interface (V747A, D748Q, W768V and C773S) also behaves in solution as a monomer. The estimated MW derived from both SEC-MALS and SAXS are matching the expected MW for a monomer and the retention times on the size exclusion column are corresponding to the retention time of the monomeric ADAR1-dsRBD3/Xlrbpa-dsRBD2 chimera and differ from the one of the wild-type ADAR1-dsRBD3, which, as already observed, elutes and behaves as a dimer in solution (Table 2 and Supplementary Figure S6). Having designed a quadruple mutant that renders ADAR1-dsRBD3 monomeric, we have now a molecular tool at hand to investigate the molecular and cellular functions of ADAR1 dimerization. In the following parts, we have used this interface mutant for exploring diverse aspects of ADAR1 biology, namely RNA-binding, cellular localization and editing activity.

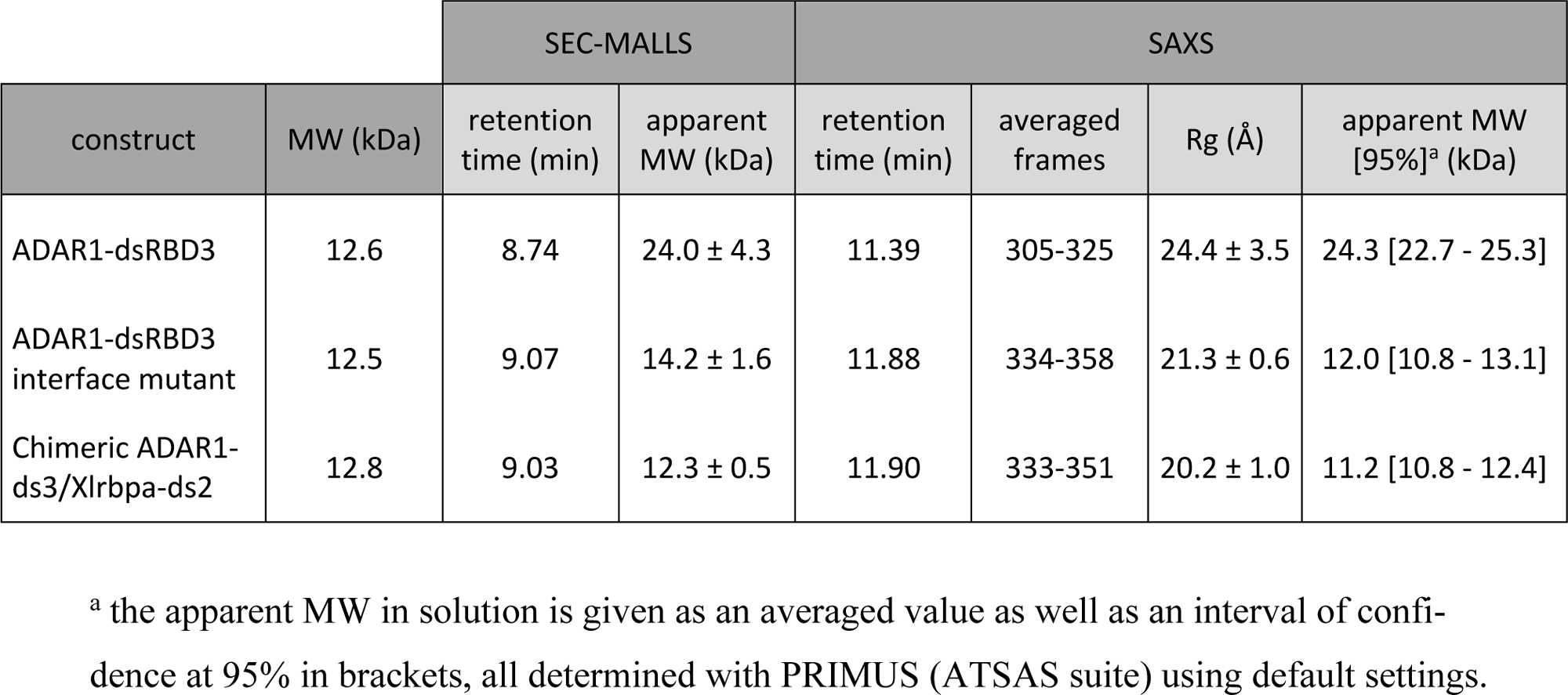
Table 2.

**Table 2**
| construct | MW (kDa) | SEC-MALLS |  | SAXS |  |  |  |
| --- | --- | --- | --- | --- | --- | --- | --- |
|  |  | retention time (min) | apparent MW (kDa) | retention time (min) | averaged frames | R<sub>g</sub> (Å) | apparent MW [95%]<sup>a</sup> (kDa) |
| ADAR1-dsRBD3 | 12.6 | 8.74 | 24.0 ± 4.3 | 11.39 | 305-325 | 24.4 ± 3.5 | 24.3 [22.7 - 25.3] |
| ADAR1-dsRBD3 interface mutant | 12.5 | 9.07 | 14.2 ± 1.6 | 11.88 | 334-358 | 21.3 ± 0.6 | 12.0 [10.8 - 13.1] |
| Chimeric ADAR1-ds3/Xlrbpa-ds2 | 12.8 | 9.03 | 12.3 ± 0.5 | 11.90 | 333-351 | 20.2 ± 1.0 | 11.2 [10.8 - 12.4] |

<sup>a</sup> the apparent MW in solution is given as an averaged value as well as an interval of confidence at 95% in brackets, all determined with PRIMUS (ATSAS suite) using default settings.

### Monomeric ADAR1-dsRBD3 remains competent for RNA-binding and nuclear localization

First, we tested whether disrupting the dimeric interface in ADAR1-dsRBD3 would drastically alter the RNA-binding capacity of the domain. For that, we measured with isothermal titration calorimetry (ITC) the dsRNA binding capacity of wild-type dimeric ADAR1-dsRBD3, monomeric ADAR1-dsRBD3 mutant, and monomeric ADAR1-dsRBD3/Xlrbpa-dsRBD2 chimera. We used an RNA duplex of 24-base pairs as a model of a dsRNA helix capable of accommodating various dsRBDs as previously described [7,46]. We showed with ITC titrations (Supplementary Figure S7), that monomeric ADAR1-dsRBD3 binds dsRNA with a strong affinity (K_D_ = 410 ± 65 nM), although its RNA-binding occurs with a slightly lower affinity compared with wild-type ADAR1-dsRBD3 (factor 3.3 ± 0.8). This corresponds to a typical RNA-binding affinity when compared to other dsRBDs [40,42,47,48], and is also similar to the binding affinity measured for the chimeric ADAR1-dsRBD3/Xlrbpa-dsRBD2 construct (K_D_ = 500 ± 55 nM – Supplementary Figure S7). Thus, disrupting dimer formation does not prevent ADAR1-dsRBD3 from binding to dsRNA.

As the bimodular NLS of ADAR1 is also flanking dsRBD3 [7], we tested whether the dimerization mutation would interfere with nuclear localization. We therefore transfected either FLAG-tagged ADAR1 p110 and ADAR1 p150 wildtype or a version harboring the dimerization mutations in dsRBD3 into HEK293 cells. Immunofluorescence staining confirmed that ADAR1 p110 remained nuclear in the presence of the dsRBD mutations preventing dimerization, while ADAR1p150 showed the expected cytoplasmic localization (Supplementary Figure S8A). Disrupting dimer formation at the level of its atypical NLS does not hinder ADAR1 from being properly delivered by the nucleocytoplasmic transport machinery.

### ADAR1-dsRBD3 mediates dimerization *in vivo*

In ADAR1, as mentioned above, dimerization had been noticed via both a region flanking dsRBD3 but also the deaminase domain itself [27,29,33]. We therefore tested the ability of dsRBD3 interface mutant for dimer formation both in the presence and absence of the deaminase domain in HEK293T cells using co-IP. To do so, wild-type and mutant versions FLAG- and HA-tagged ADAR1 variants were coexpressed in HEK293T cells. Subsequently the FLAG-tagged proteins were separated using anti-FLAG coupled magnetic beads and purified complexes were tested for the presence of HA-tagged interaction partners by western blotting. As expected, full-length versions showed an interaction irrespective of wild-type or dimerization-mutant dsRBD3, due to the presence of the deaminase domain that also presents self-interaction properties (Figure 4A) [27].

**Figure 4:**
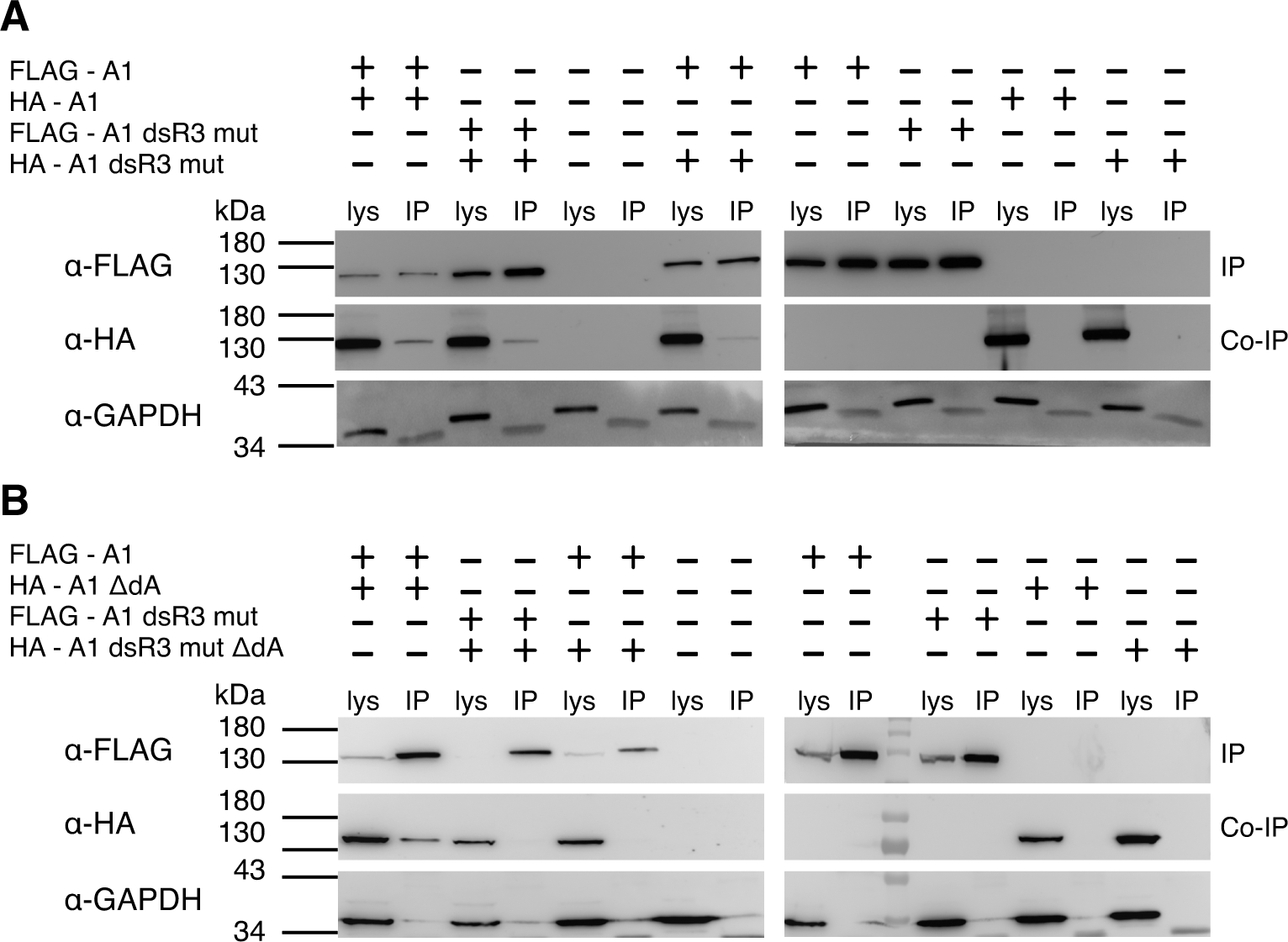
ADAR1-dsRBD3 can mediate ADAR1 dimerization *in vivo*. (**A**) Co-immunoprecipitation of FLAG-tagged and HA-tagged full length ADAR1 (FLAG-A1, HA-A1). Both protein versions were transfected as indicated by + or – signs. After precipitating the FLAG-tagged version, the precipitate was tested for the fluorescence of the HA-tagged protein in the presence of RNases. Mutations in dsRBD3 that prevent dimer formation (i.e. V747A, D748Q, W768V and C773S – FLAG/HA-A1 dsR3 mut) still allow dimer formation of full length ADAR1, likely via the deaminase domain. (**B**) In the absence of the deaminase domain (ΔdA), the dimer forming surface on dsRBD3 becomes essential for dimer formation. FLAG-tagged full length ADAR1 (FLAG-A1) interacts with HA-tagged ADAR1 without a deaminase domain (HA-A1 ΔdA). However, when the dsRBD3 is mutated to prevent dimer formation (FLAG-A1 dsR3 mut) the interaction with a deaminase deficient ADAR1 is disrupted (HA-A1 dsR3 mut ΔdA). lys: lysate; IP: immunoprecipitation; Co-IP: co-immunoprecipitation. Blots are probed with anti-FLAG (α-FLAG), anti-HA (α-HA), and anti-GAPDH (α-GAPDH) antibodies.

However, when the deaminase domain was deleted, only constructs with an intact dimerization domain in dsRBD3 could copurify, while mutations preventing dimerization of dsRBD3 disrupted dimer formation (Figure 4B). It should be noted, that all experiments were performed in the presence of RNaseA/T1 to prevent copurification of ADARs due to their joint binding to RNAs. Together, these results indicate that the dsRBD3 of ADAR1 is sufficient to mediate ADAR1 dimerization as shown in the crystal structure.

### Mutations that abolish dsRBD3 dimerization affect editing activity *in vivo*

As we had shown that both monomeric and dimeric forms of ADAR1-dsRBD3 remained competent for RNA binding *in vitro* (Figure 3 and Supplementary Figure S7), we wondered whether mutations preventing dsRBD3 dimerization would affect editing activity of either ADAR1 p150 or ADAR1 p110 *in vivo*. For this purpose, we ectopically expressed FLAG-tagged ADAR1 p150, ADAR1 p110 and the corresponding dsRBD3 interface mutants (i.e. V747A, D748Q, W768V and C773S) in HEK293T cells and measured editing levels at previously studied sites [27,49]. Expression of ADAR1 variants was tested by western blotting (Supplementary Figure S8B). To allow for the detection of site-specific editing preferences of wild-type and mutant ADAR versions, multiple sites were simultaneously probed for from one cDNA preparation. Moreover, all experiments were performed in quadruplicate and average editing levels determined.

*Azin1* carries two sites that are mostly edited by ADAR1 p150 [50,51]. Here, site 1 was edited to ∼25% by wild-type ADAR1 p150 while editing was almost completely abolished by the dimerization mutant (Figure 5A). In contrast, site 2 showed ∼10% editing by the wild-type enzyme while the dimerization mutant showed only 4% editing (Figure 5B). While this might suggest that the mutations preventing dimerization of dsRBD3 can generally reduce editing, this was not the case for a prominent site in *Gli1.* In the context of ADAR1 p150 the dimerization mutant reduced editing from ∼60% to ∼40% while the same mutation in the context of the ADAR1 p110 isoform did not lead to a significant reduction in editing (Figure 5C). To get an even better view on the impact of the dimerization mutant we analyzed editing in *Cflar*, a target with inverted SINEs and therefore multiple editing sites located in the 3’UTR. Again, editing patterns between both ADAR1 p150, ADAR1 p110 wild-type and the corresponding dsRBD3 interface mutants were investigated. Editing levels at sites 1 and 2 seemed not affected by dimerization, while editing at site 3 was strongly reduced for the mutant versions both in the context of ADAR1 p150 and ADAR1 p110 (Figure 5D, E). Sites 4, 5, and 6 are mainly targeted by ADAR1 p110. Here, a total loss of editing was seen at sites 4 and 5 for the dsRBD3 dimerization mutant in the context of ADAR1 p110 but also in the context of ADAR1 p150 (Figure 5D, E), while site 6 showed strongly reduced editing for the ADAR1 p110 dsRBD3 mutant (Figure 5E). Overall, this shows that the dimerization mutant can reduce editing at specific sites, while not affecting editing levels at other sites.

**Figure 5:**
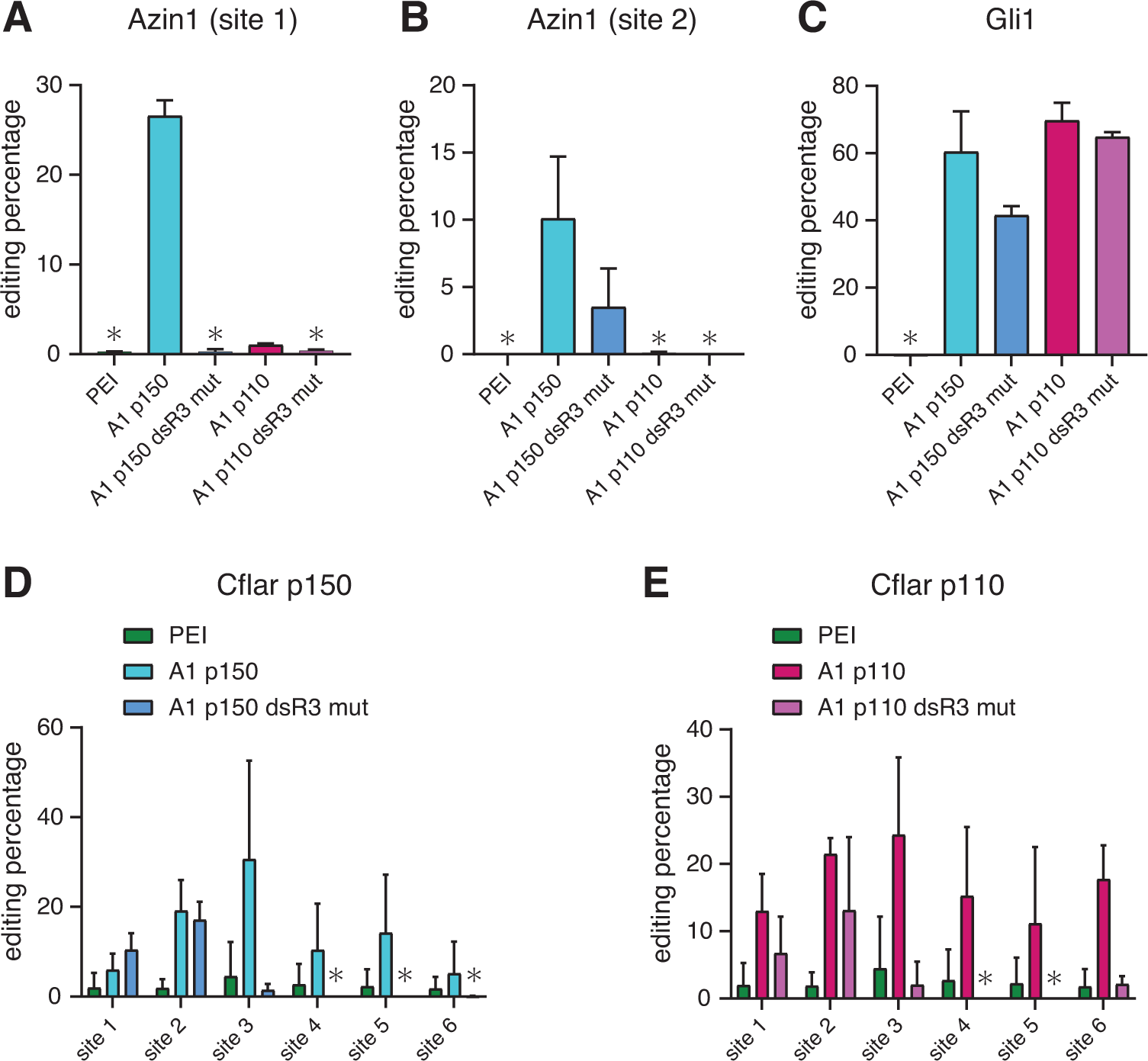
dsRBD3 dimer interface mutations affect editing activity of ADAR1 *in vivo*. HEK293T were transfected with full length ADAR1 p150 (A1 p150), ADAR1 p110 (A1 p110), ADAR1 p150 dsRBD3 mutant (A1 p150 dsR3 mut) and ADAR1 p110 dsRBD3 mutant (A1 p110 dsR3 mut). After transfection, total RNAs were extracted and different targets were amplified from cDNA. Amplicons were sequenced with Sanger sequencing and editing levels were determined by the relative height of G over A peaks. (**A**) *Azin1* site 1 is exclusively edited by ADAR1 p150 and a dimerization mutation abolishes editing. (**B**) Editing at site 2 in the same *Azin1* transcript is strongly reduced upon inhibition of dimer formation. (**C**) Editing of *Gli1* is reduced by the dimerization mutant but only in the context of ADAR1 p150. (**D**) and (**E**) Editing in the Alu element embedded within the *Cflar* transcript show that sites 1 and 2 are not affected by the loss of dimerization while editing of sites 3, 4, 5, and 6 is strongly reduced upon loss of dsRBD3 dimerization. PEI serves as a negative control, in which cells were transfected with transfection reagent only. Asterisks (*) denote no measurable editing. Error bars indicate standard deviation from 4 independent experiments.

## DISCUSSION

Here we identify a dimerization interface in ADAR1, which is located at the level of the third dsRBD of this protein. We had shown earlier that ADAR1-dsRBD3 holds an N-terminal α-helical extension (α_N_) that is important to position the N-terminal part of a bimodular nuclear localization signal that is recognized by Transportin-1 [7,52]. The crystal structure of this dsRBD could here independently confirm the N-terminal α_N_-extension, originally identified by NMR. Interestingly, however, the crystal structure of ADAR1-dsRBD3 revealed a dimerization interface, which seems unique for this particular domain as amino acids later identified to be crucial for dimerization (Supplementary Figures S4 and S6) are not found conserved on other dsRBDs [35]. The insights gained from this study have important implications for our understanding of RNA editing and will be discussed below.

Dimerization of dsRBDs had already been noticed in the dsRBD-containing proteins PACT and TRBP [53], as well as Xlrbpa, the homologue of PACT in *Xenopus laevis* [54]. In these cases, dimerization is mediated by a Type B dsRBD, namely a dsRBD that lacks specific dsRNA-binding residues and therefore cannot bind to dsRNA [32]. The homodimerization surface of these dsRBDs involve asymmetric contacts at the level of the third β-strand of the domains. The homodimer is indeed assembled through an inter-subunit parallel β-sheet where the two β3 strands interact via a register shift, thereby creating asymmetry [55]. In the case of ADAR1-dsRBD3, the dimerization reported here is clearly symmetric and consists of a back-to-back interaction of the two individual β-sheets (Figure 1), which differs from the continuous β-sheet described above. As a consequence, the symmetric homodimer of ADAR1-dsRBD3 display one set of peaks in NMR spectra (Supplementary Figure S5) [7], in contrast to PACT-dsRBD3 asymmetric homodimer that exhibits two sets of peaks [55]. Homodimerization mediated by a dsRBD has also been reported for the Staufen protein, albeit with a distinct structural organization. Again, a Type B dsRBD lacking dsRNA-binding capacities was shown to form a symmetric homodimer through a domain-swap mechanism involving an N-terminal extension to the core dsRBD structure [56]. The ADAR1-dsRBD3 dimerization reported here appears unique, since the homodimerization property is incorporated within a Type A dsRBD that has retained dsRNA-binding properties (Figure 3), while displaying protein-protein interaction capacity. This feature is encoded in the dsRBD fold itself, and not mediated by extensions attached to the dsRBD core domain, as reported previously for Type A dsRBDs [57].

The importance of ADAR dimerization for its activity has been proposed in early studies and became generally accepted in the community. In the absence of structural data supporting dimerization, this assumption was, however, based on indirect observations and mutants that did not target dimerization *per se*, but dsRNA-binding or deaminase activity of one monomer in the complex [29,31]. Recently, dimerization properties of ADARs started to be unveiled with the description of an interaction interface between deaminase domains in ADAR2 [27]. In the course of our experiments, we also noticed that the deaminase domain of ADAR1 can mediate self-association (Figure 4A), which is in good agreement with the above-mentioned study [27]. Still, our mutational analysis proved that dimerization of ADAR1 can occur in the absence of the deaminase domain in a dsRBD3-dependent manner (Figure 4B). Importantly, for the sake of comparison, interaction via the deaminase domains is not strong enough to mediate dimerization *in vitro* in the absence of dsRNA [27], whereas interaction via dsRBD3, as reported here, is strong enough to mediate dimerization even in the absence of dsRNA, and only dimeric forms of the constructs are observed *in vitro* (Figure 2 and Supplementary Figure S6). Several studies carried out on ADAR1 and/or ADAR2 pointed for an interface of interaction between the monomers to occur over a widespread region including the deaminase domain as well as the dsRNA-binding domains [29,31,33,58]. Our data, together with previous reports, therefore suggest that dsRBD3 is the main driver of dimerization in ADAR1, whereas dsRBD1 would be the equivalent region in ADAR2 [58]. On top of that, another layer of dimerization is provided by the deaminase domains, the association of which may be enhanced in the presence of dsRNA [27]. Heterodimerization of ADAR1 and ADAR2 has been observed by *in vivo* FRET experiments, and proposed as an interesting mechanism to regulate efficiency and specificity of editing [30]. Considering the absence of dsRBD3 in ADAR2, and the poor conservation of dsRBD1 between ADAR1 and ADAR2 [9], association of ADAR1 with ADAR2 is likely to occur via heterodimerization of their deaminase domains, whose interaction surfaces are conserved [27]. This suggests that assemblies composed of ADAR1 and ADAR2, if they exist at all, would be dimers of dimers bridged via their deaminase domains, an intriguing architecture that would deserve to be examined structurally.

Importantly, although dsRBD3 interface mutants retain dsRNA-binding properties, the mutations affecting dimerization have various impacts on RNA editing. For instance, the ADAR1 p150 dsRBD3 interface mutant is unable to target site 1 in *Azin1*. Similarly, dsRBD3 interface mutants of both ADAR1 p150 and p110 show altered editing patterns in the SINE containing RNA *Cflar*. However, editing of *Gli1* is unaffected in ADAR1 p110 dsRBD3 interface mutant and only slightly affected by the mutations in ADAR1 p150. Of note, mixed effects have also been observed with mutants affecting dimerization at the level of the deaminase domain in ADAR2, with editing sites that are either affected or insensitive to these mutations [27]. It is thus not straightforward to understand how dimerization affects substrate binding and site selection for editing.

We showed that dsRBD3 can bind dsRNA as a dimer (Figure 3 and Supplementary Figure S7). Given the flexible linkers between the dsRBDs of ADARs, but also between the dsRBD3 and the deaminase domain, it is difficult to predict how a substrate would be bound. While it is possible that the dsRBDs of one ADAR interact with one substrate, while the dsRBDs of the dimerization partner bind to another substrate, it is perfectly possible that the dsRBDs of both dimerized ADARs bind to the same substrate. In such a case, the region between the binding sites of one ADAR and that of the dimerization partner would need to be looped out. Double-stranded RNA is rather rigid making a looping less likely. However, single stranded RNA is very flexible and could therefore, within a few nucleotides, allow complete flexibility of two distinct double-stranded regions, such as adjacent stem-loops in the same RNA, which could be clamped by the dsRBD3 dimer.

Interestingly, our finding, that some but not all editing sites are affected when the dimerization surface of dsRBD3 is mutated suggests that dimerization is only required for efficient editing of some sites. Most importantly, this shows that dimerization has different impact on different sites, and as such, inhibition of dimerization, as opposed to inhibition of catalytic activity, might be an attractive way to pursue for modulating ADAR1 editing activity in the context of immunotherapy [26,59,60]. Towards this goal, deciphering the substrates affected by dsRBD3 dimerization will require more in-depth studies on how dimerization mutations affect transcriptome-wide editing. Positioning of affected editing sites relative to each other, would clearly shed some light on the architecture of the substrate-ADAR complexes. In this context it is interesting to note that a periodicity of editing sites has already been proposed to be affected by the binding mode of ADARs to their substrates [61,62]. In combination with the determination of the structure of full-length ADAR1 in complex with RNA substrates, such data would certainly provide some answers to the puzzling questions regarding the mechanisms of editing-site selection by ADARs.

## MATERIAL AND METHODS

### Cloning, expression, and protein purification for structural studies

The DNA sequence encoding the third dsRBD of human ADAR1 (residues 688-817: dsRBD3-long; residues 708-801: dsRBD3-mid; residues 716-797: dsRBD3-short) (Uniprot entry P55265) were subcloned by PCR amplification from full-length ADAR1, between *BamHI* and *XhoI* cloning sites in a modified pET28a expression vector containing an N-terminal tag His_6_-tag and a TEV protease cleavage site. These constructs with cleavable His_6_-tag were used in initial SAXS experiments (*i.e*. dsRBD3-long, dsRBD-mid, and dsRBD3-short) and in crystallization and structure determination (dsRBD3-short). Residues corresponding to human ADAR1-dsRDB3 (residues 708-801) were also cloned in a pET28a vector between *NdeI* and *XhoI* cloning sites. This construct with uncleavable His_6_-tag (*i.e.* ADAR1-dsRBD3), and the associated mutants generated by PCR amplifications [63] (*i.e.* ADAR1-dsRBD3 V747A+D748Q+W768V+C773S, and chimeric ADAR1-ds3/Xlrbpa-ds2), were used in SEC-MALLS, SAXS, NMR and ITC experiments. All proteins constructs were overexpressed in *E. coli* BL21(DE3) Codon-plus (RIL) cells in either LB media or M9 minimal media supplemented with ^15^NH_4_Cl. The cells were grown at 37°C to OD600 ∼0.5, cooled down at 30°C, and induced at OD600 ∼0.6-0.7 by adding isopropyl-β-D-thiogalactopyranoside to a final concentration of 1 mM. Cells were harvested 16-20 h after induction by centrifugation. Cell pellets were resuspended in lysis buffer [50 mM Tris-HCl (pH 8.0), 1 M NaCl, 20 mM imidazole, 1 mM DTT, 1mM EDTA] and lysed by sonication at 4°C. Supernatant was loaded on an Ni-NTA column on an ÄKTA purification system (Cytiva), and the protein of interest was eluted with an imidazole gradient. The fractions containing the protein were pooled and the His_6_-tag was either cleaved off by an overnight incubation with TEV protease at 20°C and removed by a second passage onto the Ni-NTA column, or kept in the final purified constructs. The proteins were further purified by size-exclusion chromatography (Superdex 75 HiLoad 26/600, Cytiva) in the storage buffer [50 mM Tris-HCl (pH 8.0), 100 mM KCl, 1mM TCEP], concentrated to ∼15-20 mg/mL using Amicon 3000 MWCO (Millipore), and kept at −20°C until further use in crystallization or biophysical assays. For specific experiments, proteins were dialyzed against the SAXS buffers [20 mM Na-HEPES pH 7.3, 55 mM KOAc, 10 mM NaCl, 1 mM TCEP] or [20 mM Na-phosphate pH 7.0, 100 mM NaCl, 2 mM 2-mercaptoethanol] or the ITC buffer [25 mM Na-phosphate pH 7.0, 100 mM NaCl, 2 mM 2-mercaptoethanol].

### SAXS data collection and analysis

SAXS experiments were performed at the SWING beamline at the SOLEIL synchrotron (Saint-Aubin, France) using an online high-performance liquid chromatography (HPLC) system equilibrated at 15°C and at a flow rate of 0.3 mL/min. All scattering intensities were collected on the elution peaks after injection on a BioSEC-3 column (Agilent) equilibrated in 20 mM Na-HEPES pH 7.3, 55 mM KOAc, 10 mM NaCl, 1 mM TCEP for the initial ADAR1-dsRBD3 constructs, or in 20 mM Na-phosphate pH 7.0, 100 mM NaCl, 2 mM 2-mercaptoethanol for the dimerization mutants. Data were processed using FOXTROT [64] and further analyzed using the ATSAS 3.0 suite of programs [65]. Data were inspected using SHANUM [66] and cropped using DATCROP [65]. The radius of gyration (Rg), the maximum dimension Dmax, as well as the molecular weight were calculated using default parameters in PRIMUS [67]. CRYSOL [68] was used to calculate theoretical scattering curves from the crystal structure atomic coordinates of dimeric and monomeric ADAR1-dsRBD3 (dsRBD-short construct). Missing side chains in the models were previously modelled in COOT assuming the most prevalent rotamer conformation.

### Crystallization, data collection, structure determination and refinement

Crystals of ADAR1-dsRBD3 (dsRBD-short construct, residues 716-797) were obtained at 5-8 mg/mL of protein by vapour diffusion against 100 mM sodium citrate pH 4.0, 20% (w/v) PEG 6000, and 1.0 M LiCl. Crystals were cryoprotected using 18-20% (v/v) glycerol before flash freezing. Data were collected on a single crystal at the micro focused PX2 beamline at the SOLEIL synchrotron (Saint-Aubin, France). Diffraction data were indexed, processed, merged and scaled using XDS [69] and AIMLESS [70]. Phases were determined by molecular replacement using PHASER [71] in the CCP4 Suite of programs [72] and the first structure of the NMR ensemble (PDB 2MDR) as a model. Model building of ADAR1-dsRBD3 was first performed with ARP/wARP using the classic warpNtrace autotracing [73]. Restrained refinements were then carried out up to 1.65 Å with the program REFMAC [74], COOT [75] and Phenix [76]. Atomic displacement parameters were modelled using a hybrid TLS + B_iso_ model, using one TLS group for each protein monomer [77].

Crystals of ADAR1-dsRBD3/dsRNA complex were obtained by mixing one equivalent of protein with 2 equivalents of RNA in 50 mM Tris-HCl pH 8.0, 100 mM KCl and 1 mM TCEP and left on ice for 1 h. The RNA sequence of 13-nucleotides employed here (*i.e.* 5’-CGAA-GCCUUCGCG-3’) has the capacity to base-pair with itself to form short double-stranded RNA modules. In addition, the nucleotide overhangs allow these RNA modules to self-assemble into a long RNA-helix, thus promoting crystal packing. Crystals of freshly prepared complex were obtained at 7 mg/mL of protein by vapour diffusion against 50 mM sodium cacodylate pH 6.0, 5% (w/v) PEG 4000, 30 mM CaCl_2_, and 230 mM KCl. Crystals were cryoprotected using 20% (v/v) glycerol before flash freezing. Data were collected on a single crystal at the micro focused PX2 beamline at the SOLEIL synchrotron (Saint-Aubin, France). Diffraction data were indexed and integrated using XDS [69] and processed using STARANISO [78] in order to take into account the moderate anisotropy of the dataset, that diffracted up to ∼2.8 Å along the best direction, but only up to ∼3.3 Å in the weakly diffracting ones. Phases were determined by molecular replacement using PHASER [71], and different combinations of models consisting of ADAR1-dsRBD3 monomer structure, double-stranded 13-nucleotides A-form RNA generated in COOT, and single-stranded 13-nucleotides adopting an A-form-like RNA structure. A complete solution (TZF = 39.0) without any missing entity in the asymmetric unit was found by searching two dsRBDs, one dsRNA module (2 x 13-nucleotides), and one ssRNA molecule (13-nucleotides). Restrained refinements were carried out up to 2.8 Å with the program LORESTR [79], REFMAC [74], and Phenix [76]. Model and map visualizations for manual reconstruction were performed with the program COOT [75]. Atomic displacement parameters were modelled using a pure TLS model, using one TLS group for each protein or RNA chain [77].

### SEC-MALLS

Size Exclusion Chromatography in line with Multi Angle Laser Light Scattering (SEC-MALLS) experiments for absolute mass determination were conducted at 25°C with a BioSEC-3 column (Agilent) equilibrated in 20 mM Na-phosphate pH 7.0, 100 mM NaCl, 2 mM 2-mercaptoethanol, on a high-performance liquid chromatography (HPLC) system (Prominence, Shimadzu) at a flow rate of 0.4 mL/min. The HPLC system is coupled to an UV detector (SPD-20, Shimadzu), an Optilab® T-rEX™ differential refractometer (Wyatt Technology) and a miniDawnTM TREOS Multi Angle Laser Light Scattering (MALLS) detector (Wyatt Technology). All data were collected and analyzed with ASTRA software (Wyatt Technology).

### NMR spectroscopy

All NMR spectra were measured at 35°C on Bruker AVIII-HD 700 MHz spectrometer equipped with a TCI 5-mm cryoprobe using 5-mm Shigemi tubes. The data were processed using Topspin 3.6 (Bruker) and analyzed with Sparky (www.cgl.ucsf.edu/home/sparky/). ADAR1-dsRBD3 mutant and chimeric proteins were checked to be properly folded by running (^15^N,^1^H)-HSQC spectra in the NMR buffer (20 mM Na-phosphate pH 7.0, 100 mM NaCl, 2 mM 2-mercaptoethanol).

### Isothermal Titration Calorimetry

To assess the nucleic acid binding properties of ADAR1-dsRBD3 interface mutant, we produced RNA substrates by in vitro transcription with T7 polymerase, namely a dsRNA duplex of 24 bp as previously described [46]. RNAs were purified by anion-exchange chromatography under native conditions as previously described [80]. DNA templates were purchased from Eurogentec. ITC experiments were performed on a VP-ITC instrument (MicroCal) calibrated according to the manufacturer’s instructions. The samples of protein and nucleic acids were prepared in and dialyzed against ITC buffer (25 mM Na-phosphate pH 7.0, 100 mM NaCl, 2 mM 2-mercaptoethanol). The concentration of protein and nucleic acid was determined using OD absorbance at 280 and 260 nm, respectively. The sample cell (1.4 mL) was loaded with 2.25 μM of the dsRNA duplex and the concentration of ADAR1-dsRBD3 wild type and variants in the syringe was 80 μM. Titration experiments were done at 25 °C with a stirring rate of 307 r.p.m. and consisted of 34 rounds of 8-μL injections. Data were plotted and analyzed using MicroCal PEAQ-ITC analysis software v1.1, using equations for a single binding-site model.

### Plasmids for HEK293T and HeLa transfections

Human ADAR1 plasmids were made as described in our previous studies [50]. All cloning and mutation procedures were done by NEBuilder HiFi DNA assembly according to the manufacturer’s instructions. All the plasmids and the oligos used for cloning and transfections are listed in Supplementary Tables S2 and S3. Western blots with FLAG or HA-tag antibodies were used for monitoring ADAR1 expression.

### Cell cultures and transfections

Henrietta Lacks (HeLa) and Human Embryonic Kidney 293 T (HEK293T) cells were cultured in high-glucose Dulbecco’s modified Eagle’s medium (DMEM) (Thermo Fischer Scientific, Waltham, MA) supplied with 10% fetal bovine serum, pyruvate and L-glutamnine. Cells were incubated at 37°C with 5% CO_2_ saturation.

Polyethyleneimine (PEI) transfections were done according to polyplus jet-PEI^R^ manufacturer’s instructions. HeLa cells were used for immunostainings and co-immunoprecipitation. HEK293T cells were used for *in vivo* editing assays.

### Co-immunoprecipitation

HEK293T cells were transfected with either FLAG-tagged ADAR1 p110 or mutant. Cells were lysed in the IP buffer [25 mM Tris-HCl pH 7.4, 150 mM NaCl, 1 mM EDTA, 1% NP-40, 5% glycerol and 1x protease inhibitor (Roche)]. The samples were lysed by BIORUPTOR® COOLER (diagenode, Belgium) using a protocol of 8 cycles of 40 seconds *on* and 60 seconds *off*, and were then treated with RNase A. The magnetic FLAG M2 beads (M8823 Merck Millipore, USA) were equilibrated in the lysis buffer and blocked with 5% BSA. The clarified clear lysate was incubated with the beads for 2 hours at 4°C on a rotating wheel. After that the beads were washed with IP buffer 5 times. The bound proteins were eluted with 2xSDS loading dye at 95°C for 5 minutes. The samples were then analysed by western blotting.

### Western blot and immunofluorescence

Ectopic expression of ADAR1 was monitored by western blotting and immunofluorescence. The cell pellet was lysed in 2xSDS sample buffer and sonicated. Lysates were separated on 8% SDS-PAGEs and transferred to Nitrocellulose membranes using a Wet-transfer. The Membranes were blocked in 1× TBST supplemented with 5% dry milk powder. ADAR constructs were detected with an anti-FLAG rabbit polyclonal antibody (Sigma, St Louis, MI), directed against the N-terminus, or an anti-HA rabbit antibody (BioLoegend, St Diego, CA). As a loading control, the lower part of the membrane was incubated with Anti-GAPDH mouse polyclonal antibody (Sigma). The antibody was diluted in 1xTBST with 3% dry milk powder. After incubation with the primary antiserum the blots were detected with a peroxidase-conjugated goat anti rabbit or mouse secondary antibody using a BioRad Chemoluminescent detection kit. Subcellular localization of ADAR1 variants were confirmed by immunofluorescent staining. In short, cells were washed with 1x PBS and fixed on coverslips using a cell-fixation solution and methanol. Then, the localization was made visible by immunofluorescence staining using an anti-flag antibody (F7425, Sigma Aldrich, Merck, Kenilworth, NJ, USA) in combination with Alexa 546 secondary antibody (Invitrogen, Thermo Fisher Scientific, Waltham, MA) and the nuclei was counterstained with DAPI. Microscopic confocal sections were taken on Confocal Laser Scanning Microscope FV3000 (Olympus, Tokyo, Japan).

### RNA isolation and editing assays

RNA isolation from the HEK293T cells was done using TRIzol^TM^ reagent (Thermo Fischer Scientific, Waltham, MA) according to manufacturer’s instructions. Isolated RNA was treated with DNaseI (New England Biolabs, Ipswich, Massachusetts) and subsequently purified by phenol:chloroform, chloroform extraction and precipitated with ethanol. The RNA was reverse transcribed using LunaScript reverse transcriptase (New England Biolabs, Ipswich, MA), random hexamer primers, Oligo(dT)s and RNase inhibitor murine. The cDNA was then used for PCR with the OneTaq 2x Master mix with Standard Buffer (New England Biolabs, Ipswich, Massachusetts). Targets for differential editing of the mutant in p150 and p110 were: *Azin1*, *Gli1* and *Cflar*. The amplicons were resolved on 2% Agarose gel to verify the size. Then a gel elution kit was used, and the samples were sent for Sanger sequencing (Eurofins, Luxembourg). The chromatograms were aligned with Geneious (Version 11.1.5) and the editing percentage was calculated with the SnapGene Viewer software. The genomic coordinates for all editing sites are provided in Supplementary Table S4.

## Author contributions

PB and MFJ conceived the study and supervised the project. AM purified constructs for crystallization studies with support from MC and with input from PB and CT; AM and PB purified constructs for SEC-MALLS and SAXS measurements; AM set-up crystallization plates and collected diffraction data; AM and PB performed diffraction data processing and refinement, SEC-MALLS and SAXS data collection and analysis; PB purified constructs, measured and analyzed NMR and ITC data. VR and SM performed *in vivo* experiments and editing assays with input from MFJ. MFJ and PB wrote the manuscript with input from AM and VR.

## Data deposition

The crystal structures of ADAR1-dsRBD3 dimer free and bound to dsRNA have been deposited to the PDB under accession numbers 7ZJ1 and 7ZLQ, respectively.

## Acknowledgments

We acknowledge SOLEIL for provision of synchrotron radiation facilities and we would like to thank William Shepard, Martin Savko and Serena Sirigu for assistance in using beamline PX2. We also thank Pierre Legrand for assistance in preliminary data collection on PX1, and for the development of the XDSME package. We are immensely indebted to Aurélien Thureau for assistance in SEC-SAXS data collection at the SWING beamline. We also acknowledge Franck Brachet for the access to the crystallography platform of the IBPC, Alexandre Pozza and Françoise Bonneté (IBPC) for assistance in SEC-MALLS data measurements and analysis, and Sylviane Hoos and Patrick England (Molecular Biophysics Facility – PFBMI, Institut Pasteur) for assistance in ITC data measurements. We also thank Alwine Hildebrandt and Tanja Rohr for excellent technical assistance. Access to the biomolecular NMR platform of the IBPC, supported by the CNRS, the Labex DYNAMO (ANR-11-LABX-0011), the Equipex CACSICE (ANR-11-EQPX-0008) and the Conseil Régional d’Île-de-France (SESAME grant) is acknowledged. PB and MFJ are supported by a joint ANR-FWF Grant (Nos. ANR-16-CE91-0003 and FWF-I2893). Part of this work was also supported by grant F80-07 by the Austrian Science Foundation.

## REFERENCES

[1] Walkley CR, Li JB (2017) Rewriting the transcriptome: adenosine-to-inosine RNA editing by ADARs. Genome Biol 18:205.

[2] Behm M, Wahlstedt H, Widmark A, Eriksson M, Öhman M (2017) Accumulation of nuclear ADAR2 regulates adenosine-to-inosine RNA editing during neuronal development. J Cell Sci 130:745–753.

[3] Roth SH, Levanon EY, Eisenberg E (2019) Genome-wide quantification of ADAR adenosine-to-inosine RNA editing activity. Nat Methods 16:1131–1138.

[4] Higuchi M, Maas S, Single FN, Hartner J, Rozov A, Burnashev N, Feldmeyer D, Sprengel R, Seeburg PH (2000) Point mutation in an AMPA receptor gene rescues lethality in mice deficient in the RNA-editing enzyme ADAR2. Nature 406:78–81.

[5] George CX, Samuel CE (1999) Human RNA-specific adenosine deaminase ADAR1 transcripts possess alternative exon 1 structures that initiate from different promoters, one constitutively active and the other interferon inducible. Proc Natl Acad Sci U S A 96:4621–6.

[6] Desterro JMP, Keegan LP, Jaffray E, Hay RT, O’Connell MA, Carmo-Fonseca M (2005) SUMO-1 modification alters ADAR1 editing activity. Mol Biol Cell 16:5115–26.

[7] Barraud P, Banerjee S, Mohamed WI, Jantsch MF, Allain FH (2014) A bimodular nuclear localization signal assembled via an extended double-stranded RNA-binding domain acts as an RNA-sensing signal for transportin 1. Proc Natl Acad Sci U S A 111:E1852–61.

[8] Poulsen H, Nilsson J, Damgaard CK, Egebjerg J, Kjems J (2001) CRM1 mediates the export of ADAR1 through a nuclear export signal within the Z-DNA binding domain. Mol Cell Biol 21:7862–71.

[9] Barraud P, Allain FH (2012) ADAR proteins: double-stranded RNA and Z-DNA binding domains. Curr Top Microbiol Immunol 353:35–60.

[10] Fritz J, Strehblow A, Taschner A, Schopoff S, Pasierbek P, Jantsch MF (2009) RNA-regulated interaction of transportin-1 and exportin-5 with the double-stranded RNA-binding domain regulates nucleocytoplasmic shuttling of ADAR1. Mol Cell Biol 29:1487–97.

[11] Licht K, Jantsch MF (2017) The Other Face of an Editor: ADAR1 Functions in Editing-Independent Ways. Bioessays 39:11.

[12] Kim JI, Nakahama T, Yamasaki R, Costa Cruz PH, Vongpipatana T, Inoue M, Kanou N, Xing Y, Todo H, Shibuya T, Kato Y, Kawahara Y (2021) RNA editing at a limited number of sites is sufficient to prevent MDA5 activation in the mouse brain. PLoS Genet 17:e1009516.

[13] Dorrity TJ, Chung H (2023) A tale of two pathways: Two distinct mechanisms of ADAR1 prevent fatal autoinflammation. Mol Cell 83:3760–3762.

[14] Liddicoat BJ, Piskol R, Chalk AM, Ramaswami G, Higuchi M, Hartner JC, Li JB, Seeburg PH, Walkley CR (2015) RNA editing by ADAR1 prevents MDA5 sensing of endogenous dsRNA as nonself. Science 349:1115–20.

[15] Mannion NM, Greenwood SM, Young R, Cox S, Brindle J, Read D, Nellåker C, Vesely C, Ponting CP, McLaughlin PJ, Jantsch MF, Dorin J, Adams IR, Scadden ADJ, Ohman M, Keegan LP, O’Connell MA (2014) The RNA-editing enzyme ADAR1 controls innate immune responses to RNA. Cell Rep 9:1482–94.

[16] Pestal K, Funk CC, Snyder JM, Price ND, Treuting PM, Stetson DB (2015) Isoforms of RNA-Editing Enzyme ADAR1 Independently Control Nucleic Acid Sensor MDA5-Driven Autoimmunity and Multi-organ Development. Immunity 43:933–44.

[17] Chung H, Calis JJA, Wu X, Sun T, Yu Y, Sarbanes SL, Dao Thi VL, Shilvock AR, Hoffmann H, Rosenberg BR, Rice CM (2018) Human ADAR1 Prevents Endogenous RNA from Triggering Translational Shutdown. Cell 172:811–824.

[18] Hu S, Heraud-Farlow J, Sun T, Liang Z, Goradia A, Taylor S, Walkley CR, Li JB (2023) ADAR1p150 prevents MDA5 and PKR activation via distinct mechanisms to avert fatal auto-inflammation. Mol Cell 83:3869–3884.

[19] de Reuver R, Verdonck S, Dierick E, Nemegeer J, Hessmann E, Ahmad S, Jans M, Blancke G, Van Nieuwerburgh F, Botzki A, Vereecke L, van Loo G, Declercq W, Hur S, Vandenabeele P, Maelfait J (2022) ADAR1 prevents autoinflammation by suppressing spontaneous ZBP1 activation. Nature 607:784–789.

[20] Hubbard NW, Ames JM, Maurano M, Chu LH, Somfleth KY, Gokhale NS, Werner M, Snyder JM, Lichauco K, Savan R, Stetson DB, Oberst A (2022) ADAR1 mutation causes ZBP1-dependent immunopathology. Nature 607:769–775.

[21] Zhang T, Yin C, Fedorov A, Qiao L, Bao H, Beknazarov N, Wang S, Gautam A, Williams RM, Crawford JC, Peri S, Studitsky V, Beg AA, Thomas PG, Walkley C, Xu Y, Poptsova M, Herbert A, Balachandran S (2022) ADAR1 masks the cancer immunotherapeutic promise of ZBP1-driven necroptosis. Nature 606:594–602.

[22] Haudenschild BL, Maydanovych O, Véliz EA, Macbeth MR, Bass BL, Beal PA (2004) A transition state analogue for an RNA-editing reaction. J Am Chem Soc 126:11213–9.

[23] Eggington JM, Greene T, Bass BL (2011) Predicting sites of ADAR editing in double-stranded RNA. Nat Commun 2:319.

[24] Nishikura K (2010) Functions and regulation of RNA editing by ADAR deaminases. Annu Rev Biochem 79:321–49.

[25] Mannion N, Arieti F, Gallo A, Keegan LP, O’Connell MA (2015) New Insights into the Biological Role of Mammalian ADARs; the RNA Editing Proteins. Biomolecules 5:2338–62.

[26] Pecori R, Papavasiliou NF (2020) It takes two (and some distance) to tango: how ADARs join to edit RNA. Nat Struct Mol Biol 27:308–310.

[27] Thuy-Boun AS, Thomas JM, Grajo HL, Palumbo CM, Park S, Nguyen LT, Fisher AJ, Beal PA (2020) Asymmetric dimerization of adenosine deaminase acting on RNA facilitates substrate recognition. Nucleic Acids Res 48:7958–7972.

[28] Gallo A, Keegan LP, Ring GM, O’Connell MA (2003) An ADAR that edits transcripts encoding ion channel subunits functions as a dimer. EMBO J 22:3421–30.

[29] Cho DC, Yang W, Lee JT, Shiekhattar R, Murray JM, Nishikura K (2003) Requirement of dimerization for RNA editing activity of adenosine deaminases acting on RNA. J Biol Chem 278:17093–102.

[30] Chilibeck KA, Wu T, Liang C, Schellenberg MJ, Gesner EM, Lynch JM, MacMillan AM (2006) FRET analysis of in vivo dimerization by RNA-editing enzymes. J Biol Chem 281:16530–5.

[31] Valente L, Nishikura K (2007) RNA binding-independent dimerization of adenosine deaminases acting on RNA and dominant negative effects of nonfunctional subunits on dimer functions. J Biol Chem 282:16054–61.

[32] Gleghorn ML, Maquat LE (2014) ’Black sheep’ that don’t leave the double-stranded RNA-binding domain fold. Trends Biochem Sci 39:328–40.

[33] Ota H, Sakurai M, Gupta R, Valente L, Wulff B, Ariyoshi K, Iizasa H, Davuluri RV, Nishikura K (2013) ADAR1 forms a complex with Dicer to promote microRNA processing and RNA-induced gene silencing. Cell 153:575–89.

[34] Bou-Nader C, Barraud P, Pecqueur L, Pérez J, Velours C, Shepard W, Fontecave M, Tisné C, Hamdane D (2019) Molecular basis for transfer RNA recognition by the double-stranded RNA-binding domain of human dihydrouridine synthase 2. Nucleic Acids Res 47:3117–3126.

[35] Masliah G, Barraud P, Allain FH (2013) RNA recognition by double-stranded RNA binding domains: a matter of shape and sequence. Cell Mol Life Sci 70:1875–95.

[36] Ryter JM, Schultz SC (1998) Molecular basis of double-stranded RNA-protein interactions: structure of a dsRNA-binding domain complexed with dsRNA. EMBO J 17:7505–13.

[37] Ramos A, Grünert S, Adams J, Micklem DR, Proctor MR, Freund S, Bycroft M, St Johnston D, Varani G (2000) RNA recognition by a Staufen double-stranded RNA-binding domain. EMBO J 19:997–1009.

[38] Wu H, Henras A, Chanfreau G, Feigon J (2004) Structural basis for recognition of the AGNN tetraloop RNA fold by the double-stranded RNA-binding domain of Rnt1p RNase III. Proc Natl Acad Sci U S A 101:8307–12.

[39] Gan J, Tropea JE, Austin BP, Court DL, Waugh DS, Ji X (2006) Structural insight into the mechanism of double-stranded RNA processing by ribonuclease III. Cell 124:355–66.

[40] Stefl R, Oberstrass FC, Hood JL, Jourdan M, Zimmermann M, Skrisovska L, Maris C, Peng L, Hofr C, Emeson RB, Allain FH (2010) The solution structure of the ADAR2 dsRBM-RNA complex reveals a sequence-specific readout of the minor groove. Cell 143:225–37.

[41] Jayachandran U, Grey H, Cook AG (2016) Nuclear factor 90 uses an ADAR2-like binding mode to recognize specific bases in dsRNA. Nucleic Acids Res 44:1924–36.

[42] Lazzaretti D, Bandholz-Cajamarca L, Emmerich C, Schaaf K, Basquin C, Irion U, Bono F (2018) The crystal structure of Staufen1 in complex with a physiological RNA sheds light on substrate selectivity. Life Sci Alliance 1:e201800187.

[43] Yadav DK, Zigáčková D, Zlobina M, Klumpler T, Beaumont C, Kubíčková M, Vaňáčová Š, Lukavsky PJ (2020) Staufen1 reads out structure and sequence features in ARF1 dsRNA for target recognition. Nucleic Acids Res 48:2091–2106.

[44] Jin W, Wang J, Liu C, Wang H, Xu R (2020) Structural Basis for pri-miRNA Recognition by Drosha. Mol Cell 78:423–433.

[45] Castrignanò T, Chillemi G, Varani G, Desideri A (2002) Molecular dynamics simulation of the RNA complex of a double-stranded RNA-binding domain reveals dynamic features of the intermolecular interface and its hydration. Biophys J 83:3542–52.

[46] Barraud P, Emmerth S, Shimada Y, Hotz H, Allain FH, Bühler M (2011) An extended dsRBD with a novel zinc-binding motif mediates nuclear retention of fission yeast Dicer. EMBO J 30:4223–35.

[47] Barraud P, Heale BSE, O’Connell MA, Allain FH (2012) Solution structure of the N-terminal dsRBD of Drosophila ADAR and interaction studies with RNA. Biochimie 94:1499–509.

[48] Yamashita S, Nagata T, Kawazoe M, Takemoto C, Kigawa T, Güntert P, Kobayashi N, Terada T, Shirouzu M, Wakiyama M, Muto Y, Yokoyama S (2011) Structures of the first and second double-stranded RNA-binding domains of human TAR RNA-binding protein. Protein Sci 20:118–30.

[49] Levanon EY, Eisenberg E, Yelin R, Nemzer S, Hallegger M, Shemesh R, Fligelman ZY, Shoshan A, Pollock SR, Sztybel D, Olshansky M, Rechavi G, Jantsch MF (2004) Systematic identification of abundant A-to-I editing sites in the human transcriptome. Nat Biotechnol 22:1001–5.

[50] Kleinova R, Rajendra V, Leuchtenberger AF, Lo Giudice C, Vesely C, Kapoor U, Tanzer A, Derdak S, Picardi E, Jantsch MF (2023) The ADAR1 editome reveals drivers of editing-specificity for ADAR1-isoforms. Nucleic Acids Res 51:4191–4207.

[51] Xing Y, Nakahama T, Wu Y, Inoue M, Kim JI, Todo H, Shibuya T, Kato Y, Kawahara Y (2023) RNA editing of AZIN1 coding sites is catalyzed by ADAR1 p150 after splicing. J Biol Chem 299:104840.

[52] Mboukou A, Rajendra V, Kleinova R, Tisné C, Jantsch MF, Barraud P (2021) Transportin-1: A Nuclear Import Receptor with Moonlighting Functions. Front Mol Biosci 8:638149.

[53] Laraki G, Clerzius G, Daher A, Melendez-Peña C, Daniels S, Gatignol A (2008) Interactions between the double-stranded RNA-binding proteins TRBP and PACT define the Medipal domain that mediates protein-protein interactions. RNA Biol 5:92–103.

[54] Hitti EG, Sallacz NB, Schoft VK, Jantsch MF (2004) Oligomerization activity of a double-stranded RNA-binding domain. FEBS Lett 574:25–30.

[55] Heyam A, Coupland CE, Dégut C, Haley RA, Baxter NJ, Jakob L, Aguiar PM, Meister G, Williamson MP, Lagos D, Plevin MJ (2017) Conserved asymmetry underpins homodimerization of Dicer-associated double-stranded RNA-binding proteins. Nucleic Acids Res 45:12577–12584.

[56] Gleghorn ML, Gong C, Kielkopf CL, Maquat LE (2013) Staufen1 dimerizes through a conserved motif and a degenerate dsRNA-binding domain to promote mRNA decay. Nat Struct Mol Biol 20:515–24.

[57] Banerjee S, Barraud P (2014) Functions of double-stranded RNA-binding domains in nucleocytoplasmic transport. RNA Biol 11:1226–32.

[58] Poulsen H, Jorgensen R, Heding A, Nielsen FC, Bonven B, Egebjerg J (2006) Dimerization of ADAR2 is mediated by the double-stranded RNA binding domain. RNA 12:1350–60.

[59] Ishizuka JJ, Manguso RT, Cheruiyot CK, Bi K, Panda A, Iracheta-Vellve A, Miller BC, Du PP, Yates KB, Dubrot J, Buchumenski I, Comstock DE, Brown FD, Ayer A, Kohnle IC, Pope HW, Zimmer MD, Sen DR, Lane-Reticker SK, Robitschek EJ, Griffin GK, Collins NB, Long AH, Doench JG, Kozono D, Levanon EY, Haining WN (2019) Loss of ADAR1 in tumours overcomes resistance to immune checkpoint blockade. Nature 565:43–48.

[60] Datta R, Adamska JZ, Bhate A, Li JB (2023) A-to-I RNA editing by ADAR and its therapeutic applications: From viral infections to cancer immunotherapy. Wiley Interdiscip Rev RNA in press:e1817.

[61] Song Y, Yang W, Fu Q, Wu L, Zhao X, Zhang Y, Zhang R (2020) irCLASH reveals RNA substrates recognized by human ADARs. Nat Struct Mol Biol 27:351–362.

[62] Uzonyi A, Nir R, Shliefer O, Stern-Ginossar N, Antebi Y, Stelzer Y, Levanon EY, Schwartz S (2021) Deciphering the principles of the RNA editing code via large-scale systematic probing. Mol Cell 81:2374–2387.

[63] Liu H, Naismith JH (2008) An efficient one-step site-directed deletion, insertion, single and multiple-site plasmid mutagenesis protocol. BMC Biotechnol 8:91.

[64] Thureau A, Roblin P, Pérez J (2021) BioSAXS on the SWING beamline at Synchrotron SOLEIL. Journal of Applied Crystallography 54:1698–1710.

[65] Manalastas-Cantos K, Konarev PV, Hajizadeh NR, Kikhney AG, Petoukhov MV, Molodenskiy DS, Panjkovich A, Mertens HDT, Gruzinov A, Borges C, Jeffries CM, Svergun DI, Franke D (2021) ATSAS 3.0: expanded functionality and new tools for small-angle scattering data analysis. Journal of Applied Crystallography 54:343–355.

[66] Konarev PV, Svergun DI (2015) A posteriori determination of the useful data range for small-angle scattering experiments on dilute monodisperse systems. IUCrJ 2:352–360.

[67] Konarev PV, Volkov VV, Sokolova AV, Koch MHJ, Svergun DI (2003) PRIMUS: a Windows PC-based system for small-angle scattering data analysis. Journal of Applied Crystallography 36:1277–1282.

[68] Svergun D, Barberato C, Koch MHJ (1995) CRYSOL - a Program to Evaluate X-ray Solution Scattering of Biological Macromolecules from Atomic Coordinates. Journal of Applied Crystallography 28:768–773.

[69] Kabsch W (2010) XDS. Acta Crystallogr D Biol Crystallogr 66:125–32.

[70] Evans PR, Murshudov GN (2013) How good are my data and what is the resolution? Acta Crystallogr D Biol Crystallogr 69:1204–14.

[71] McCoy AJ, Grosse-Kunstleve RW, Adams PD, Winn MD, Storoni LC, Read RJ (2007) Phaser crystallographic software. J Appl Crystallogr 40:658–674.

[72] Winn MD, Ballard CC, Cowtan KD, Dodson EJ, Emsley P, Evans PR, Keegan RM, Krissinel EB, Leslie AGW, McCoy A, McNicholas SJ, Murshudov GN, Pannu NS, Potterton EA, Powell HR, Read RJ, Vagin A, Wilson KS (2011) Overview of the CCP4 suite and current developments. Acta Crystallogr D Biol Crystallogr 67:235–42.

[73] Langer G, Cohen SX, Lamzin VS, Perrakis A (2008) Automated macromolecular model building for X-ray crystallography using ARP/wARP version 7. Nat Protoc 3:1171–9.

[74] Murshudov GN, Skubák P, Lebedev AA, Pannu NS, Steiner RA, Nicholls RA, Winn MD, Long F, Vagin AA (2011) REFMAC5 for the refinement of macromolecular crystal structures. Acta Crystallogr D Biol Crystallogr 67:355–67.

[75] Emsley P, Lohkamp B, Scott WG, Cowtan K (2010) Features and development of Coot. Acta Crystallogr D Biol Crystallogr 66:486–501.

[76] Liebschner D, Afonine PV, Baker ML, Bunkóczi G, Chen VB, Croll TI, Hintze B, Hung LW, Jain S, McCoy AJ, Moriarty NW, Oeffner RD, Poon BK, Prisant MG, Read RJ, Richardson JS, Richardson DC, Sammito MD, Sobolev OV, Stockwell DH, Terwilliger TC, Urzhumtsev AG, Videau LL, Williams CJ, Adams PD (2019) Macromolecular structure determination using X-rays, neutrons and electrons: recent developments in Phenix. Acta Crystallogr D Struct Biol 75:861–877.

[77] Merritt EA (2012) To B or not to B: a question of resolution? Acta Crystallogr D Biol Crystallogr 68:468–77.

[78] Vonrhein C, Tickle IJ, Flensburg C, Keller P, Paciorek W, Sharff A, Bricogne G (2018) Advances in automated data analysis and processing within autoPROC, combined with improved characterisation, mitigation and visualisation of the anisotropy of diffraction limits using STARANISO. Acta Crystallogr. A 74:A360.

[79] Kovalevskiy O, Nicholls RA, Murshudov GN (2016) Automated refinement of macromolecular structures at low resolution using prior information. Acta Crystallogr D Struct Biol 72:1149–1161.

[80] Catala M, Gato A, Tisné C, Barraud P (2020) Preparation of Yeast tRNA Sample for NMR Spectroscopy. Bio Protoc 10:e3646.

