## Supplementary information for "Dimerization of ADAR1 modulates site-specificity of RNA editing"

**Content:**

Supplementary Table S1 to S4 page 2-4
Supplementary Figures S1 to S8 pages 5-12

**Supplementary Table S1: Data collection and refinement statistics.**

| Structure | ADAR1-dsRBD3 dimer | ADAR1-dsRBD3:dsRNA |
|---|---|---|
| **PDB code** | 7ZJ1 | 7ZLQ |
| **Data collection** | | |
| **Beamline** | SOLEIL Proxima-2 (PX2) | SOLEIL Proxima-2 (PX2) |
| **Wavelength** | 0.9801 | 0.9999 |
| **Space group** | P 31 2 1 | P 61 2 2 |
| **Unit cell** | 44.2353 44.2353 131.803 90 90 120 | 91.294 91.294 207.527 90 90 120 |
| **Resolution range** | 19.06–1.65 (1.71–1.65) | 44.58–2.80 (2.99–2.80) |
| **Total reflections** | 351229 (26321) | 192634 (9181) |
| **Unique reflections** | 18792 (1852) | 10044 (503) |
| **Multiplicity** | 18.7 (14.2) | 19.2 (18.3) |
| **Completeness (%)** | 99.26 (99.84) | 75.9 (22.1) [94.9 (88.6)][a] |
| **Mean I/sigma(I)** | 12.45 (1.53) | 16.6 (1.2) |
| **R-merge** | 0.1518 (0.8674) | 0.163 (2.663) |
| **R-pim** | 0.0354 (0.2381) | 0.038 (0.633) |
| **$CC_{1/2}$** | 0.995 (0.585) | 1.000 (0.517) |
| **Refinement** | | |
| **Reflections used in refinement** | 18670 (1854) | 10026 (503) |
| **Reflections used for R-free** | 943 (92) | 1078 (73) |
| **R-work** | 0.1871 (0.2079) | 0.2311 (0.3278) |
| **R-free** | 0.2247 (0.2540) | 0.2658 (0.3526) |
| **Model composition** | | |
| **Number of non-hydrogen atoms** | 1364 | 2079 |
| **macromolecules** | 1248 | 2079 |
| **solvent** | 116 | 0 |
| **Protein residues** | 166 | 164 |
| **RNA residues** | 0 | 39 |
| **Model validation** | | |
| **Clashscore** | 3.25 | 7.99 |
| **Rotamer outliers (%)** | 0.00 | 0.00 |
| **Average B-factor (Å$^2$)** | 20.45 | 80.97 |
| **protein** | 19.87 | 79.95 |
| **RNA** | – | 82.54 |
| **solvent** | 26.67 | – |
| **Number of TLS groups** | 2 | 5 |
| **Ramachandran statistics (%)** | | |
| **Favored** | 98.15 | 97.50 |
| **Allowed** | 1.85 | 2.50 |
| **Outliers** | 0.00 | 0.00 |
| **RMS deviations** | | |
| **Bond length (Å)** | 0.010 | 0.007 |
| **Bond angles (°)** | 1.08 | 0.88 |

Statistics for the highest-resolution shell are shown in parentheses.
[a]Values in brackets correspond to the ellipsoidal completeness as reported by STARANISO

**Supplementary Table S2: Oligos**

| | | |
|---|---|---|
| MJ8371 | GATACCTGAACACCAACCCTGTGGGTGGCCTTTTGGAGTAC | Insert forward primer used for creating plasmids MJ1645,1648 |
| MJ8372 | CCCCAATCAAGACACGGAGAGCCGCATCTGCTGCTTCCTG | Insert reverse primer used for creating plasmids MJ1645,1648 |
| MJ8373 | GATGCGGCTCTCCGTGTCTTGATTGGGGAGAACGAGAAGG | Vector forward primer used for creating plasmids MJ1645,1648 |
| MJ8374 | CCACCCACAGGGTTGGTGTTCAGGTATCTCACGAGCTCGCC | Vector reverse primer used for creating plasmids MJ1645,1648 |
| MJ8610 | GGCCACCATGTACCCATACGATGTTCCAGATTACGCTATGGCCGAGATCAAGG | Insert forward primer used for creating plasmids MJ1723, MJ1724 |
| MJ8611 | CCCTCTCCACTGCCGACTAGTACTGGGCAGAGATAAAAGTTCTTTTCCTCCTG | Insert reverse primer used for creating plasmids MJ1723, MJ1724 |
| MJ8612 | CTGCCCAGTACTAGTCGGCAGTGGAGAGGGCAGAGGAAGTCTGCTAACATG | Vector forward primer used for creating plasmids MJ1723, MJ1724 |
| MJ8613 | GCGTAATCTGGAACATCGTATGGGTACATGGTGGCCAGATATCCAGCACAG | Vector reverse primer used for creating plasmids MJ1723, MJ1724 |
| MJ8697 | GCAGTGGAGAGGGCAGAGGAAGTCTGCTAACATGCGGTG | Vector forward primer used for creating plasmidsMJ1731, MJ1732 |
| MJ8698 | GCAGACTTCCTCTGCCCTCTCCACTGCCGAGTGTCTTTGGCTG | Insert reverse primer used for creating plasmidsMJ1731, MJ1732 |
| MJ8749 | GAAGGATCTGGTGTTAAGATAATTTCAGAACCCG | Forward primer Azin1 |
| MJ8750 | ACTGGAATGTTGACCAGACAAGCTTAACC | Reverse primer Azin1 |
| MJ8755 | CGAGCCGAGTATCCAGGATACAAC | Forward primer Gli1 |
| MJ8756 | CCCATATCCCAGAGTATCAGTAGGTGG | Reverse primer Gli1 |
| MJ984 | CCCACTCCTGGATCTTCAC | Forward primer Cflar |
| MJ985 | CAGGTTGGTATGCAGTGGC | Reverse primer Cflar |

* MJ= oligo numbers according Jantsch lab oligo database

**Supplementary Table S3: Mammalian expression plasmids.**

| | |
|---|---|
| MJ1209 | FLAG-human ADAR1 p110-6X His in pCDNA 3.1, Gift from Mary O' Connell lab |
| MJ1210 | FLAG-human ADAR1 p150-6X His in pCDNA 3.1, Gift from Mary O' Connell lab |
| MJ1645 | FLAG-human ADAR1 p110 dsRBD3 mutant-6X His in pCDNA 3.1 |
| MJ1648 | FLAG-human ADAR1 p150 dsRBD3 mutant-6X His in pCDNA 3.1 |
| MJ1723 | HA-human ADAR1 p110 T2A eGFP in pCDNA 3.1 |
| MJ1724 | HA-human ADAR1 p110 dsRBD3 mutant T2A eGFP in pCDNA 3.1 |
| MJ1731 | HA-human ADAR1 p110 Δ deaminase T2A eGFP in pCDNA 3.1 |
| MJ1732 | HA-human ADAR1 p110 dsRBD3 mutant Δ deaminase T2A eGFP in pCDNA 3.1 |

* MJ= plasmid numbers according Jantsch lab plasmid database, MJ1731 and MJ1732 expresses amino acids from 296 to 833 (Methionine 296 is the start for p110 variant of ADAR1)

**Supplementary Table S4: Genomic coordinates of editing sites.**

| | |
|---|---|
| Cflar Site 1 | chr2:201164032 |
| Cflar Site 2 | chr2:201164087 |
| Cflar Site 3 | chr2:201164088 |
| Cflar Site 4 | chr2:201164105 |
| Cflar Site 5 | chr2:201164112 |
| Cflar Site 6 | chr2:201164118 |
| Azin1 Site 1 | chr8:102829408 |
| Azin1 Site 2 | chr8:102829879 |
| Gli | chr12:57470841 |

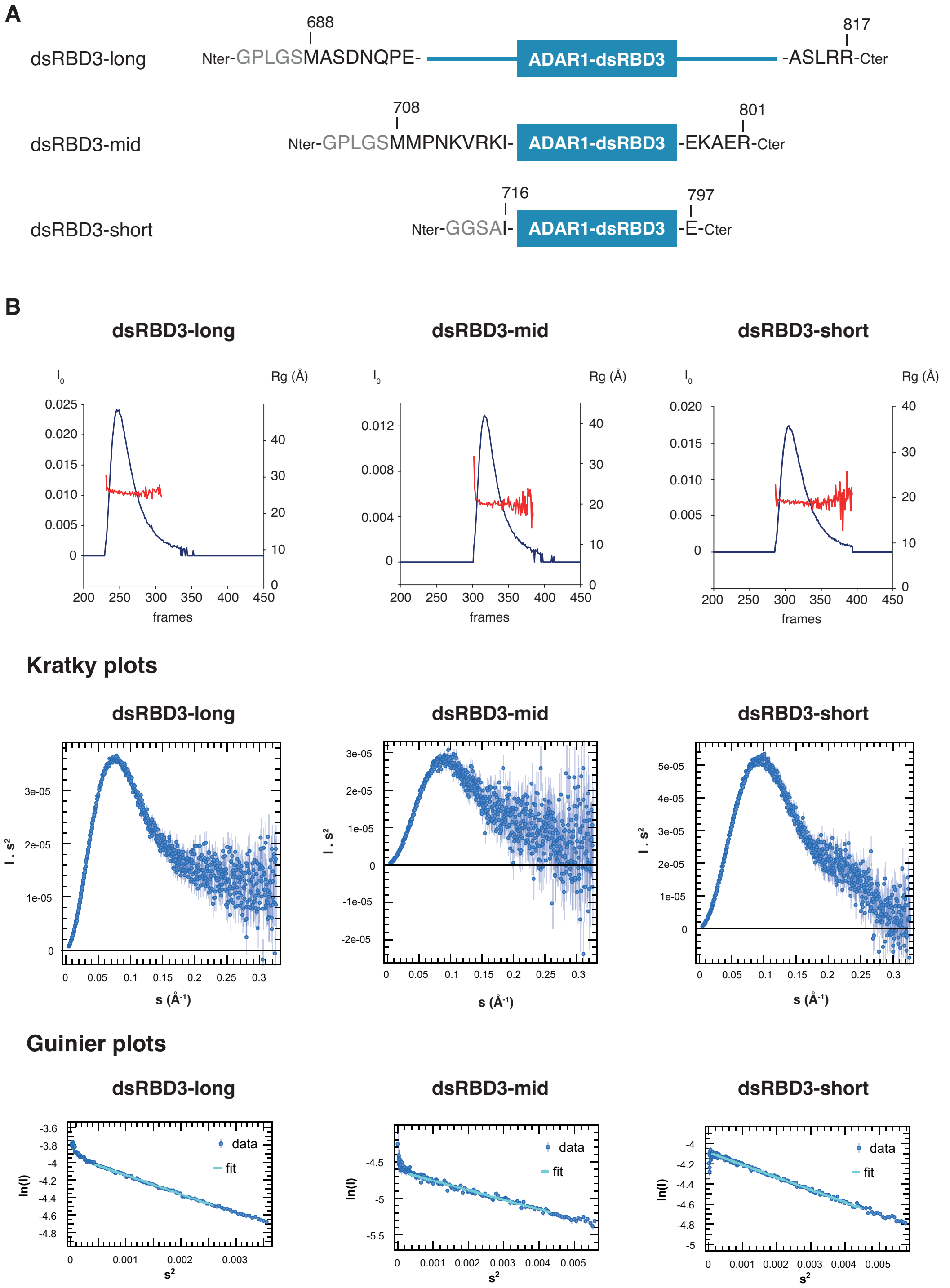

**Supplementary Figure S1: Characterization of various ADAR1-dsRBD3 constructs with different length of N- and C-terminal flanking tails by SAXS.**

(**A**) Schematic representation of ADAR1-dsRBD3 constructs: dsRBD3-long (residues 688-817), dsRBD3-mid (residues 708-801) and dsRBD3-short (residues 716-797). Constructs used here had their purification tags cleaved off. Remaining residues after tag-cleavage are shown in grey. (**B**) SAXS data are presented for the ADAR1-dsRBD3 constructs displayed on panel A. $I_0$ and Rg values obtained around the elution peak of HPLC are displayed in the upper panel. Kratky plots are shown in the middle panel (Kratky plots). Regions used for the Guinier approximation are shown in the lower panel (Guinier plots). Derived parameters are reported on Table 1.

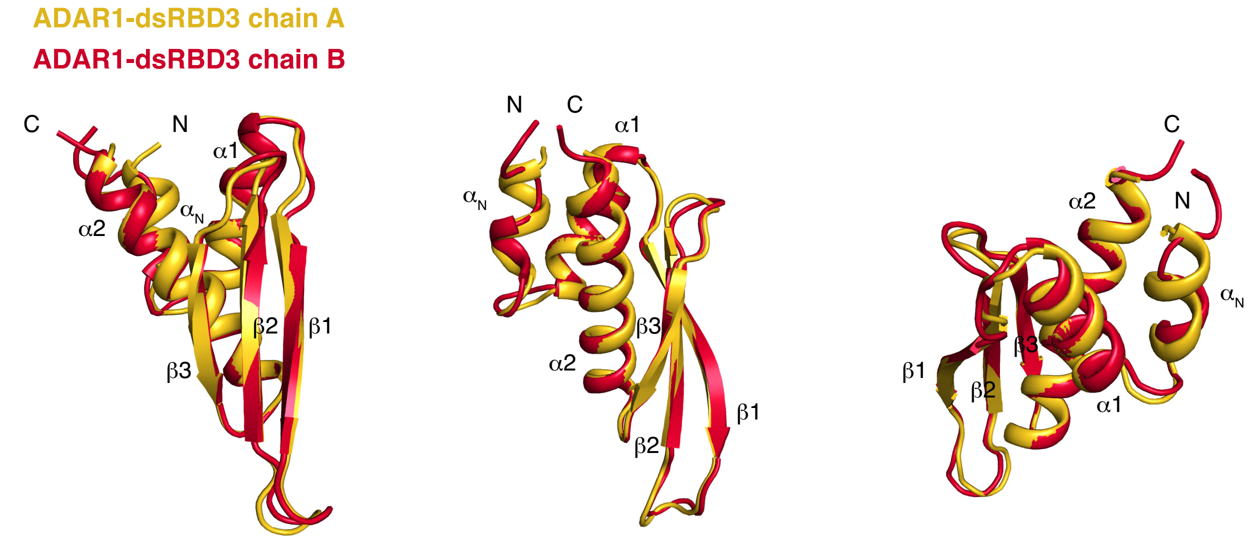

**Supplementary Figure S2: Comparison of ADAR1-dsRBD3 chain A and chain B in the asymmetric unit.**

ADAR1-dsRBD3 chain A (*in yellow*) and chain B (*in red*) are shown as cartoon. Both chains were superimposed over Cα atoms over the entire domain. Secondary structure elements are labelled. Most differences occur at the level of the N-terminal helix $\alpha_N$, where the distortion on chain B is likely caused by packing interactions with symmetrical molecules. Small structural differences also occur at the level of loop 1 (α1-β1), loop 2 (β1-β2), and loop 3 (β2-β3).

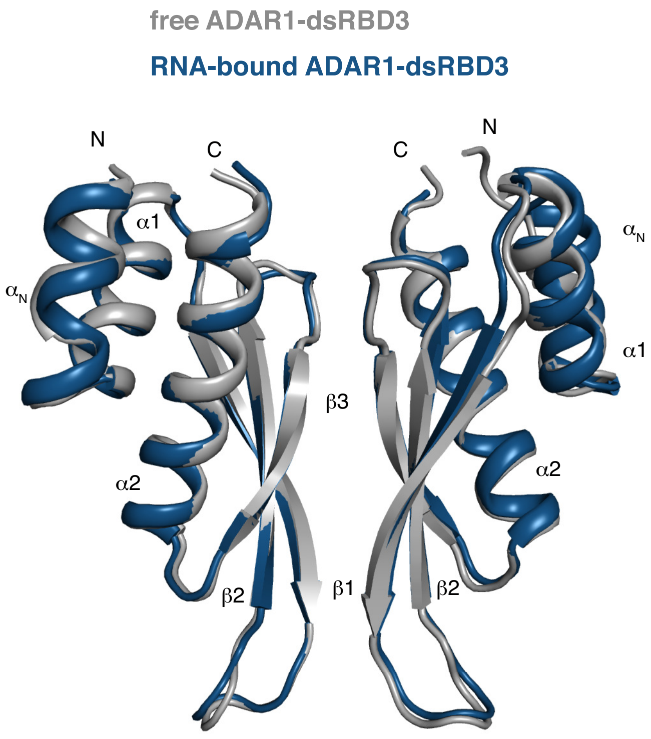

**Supplementary Figure S3: Comparison of free ADAR1-dsRBD3 dimer and RNA-bound ADAR1-dsRBD3 dimer.**

Free ADAR1-dsRBD3 (*in grey*) and RNA-bound ADAR1-dsRBD3 (*in blue*) are shown as cartoon. Both structures were superimposed over Cα atoms over the entire domains. Secondary structure elements are labelled. The two structures are perfectly superimposable, meaning that the dimer organization is identical in both structures.

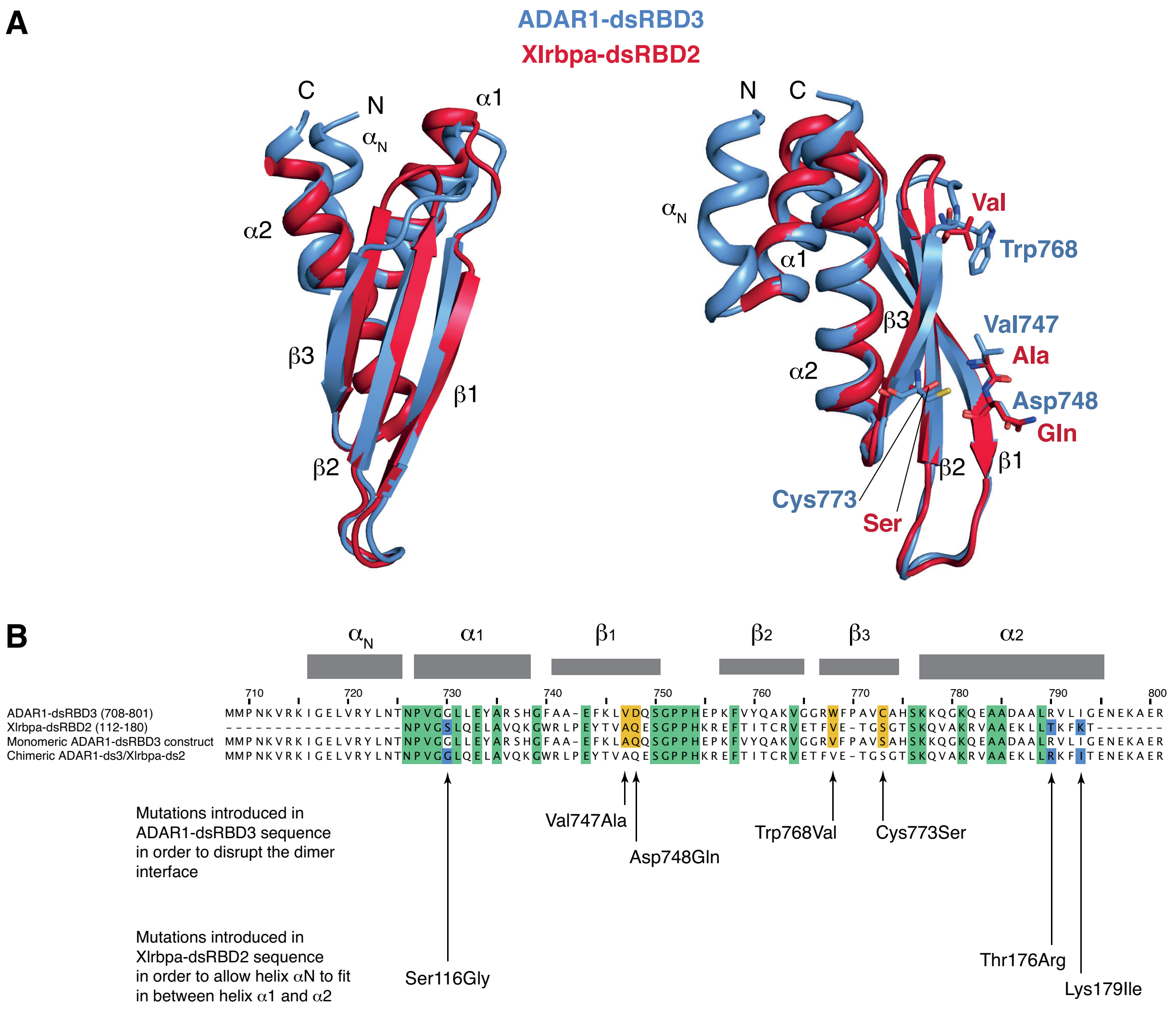

**Supplementary Figure S4: Rational design of ADAR1-dsRBD3 mutants to disrupt the dimer interface.**

(**A**) ADAR1-dsRBD3 (*in light blue*) and Xlrbpa-dsRBD2 (*in red*) are shown as cartoon. Both chains were superimposed over Cα atoms over the entire domains. Secondary structure elements are labelled. Residues at the β-sheet interface involved in the monomer-monomer interaction (see Figure 1) and that differ between ADAR1-dsRBD3 and Xlrbpa-dsRBD2 (see the sequence alignment on panel B), are shown as sticks. (**B**) Sequence alignment of ADAR1-dsRBD3 (708-801), Xlrbpa-dsRBD2 (112-180), the monomeric ADAR1-dsRBD3 construct that include four point-mutations at the dimer interface in which ADAR1-dsRBD3 residues are mutated into those found in Xlrbpa-dsRBD2 (i.e. V747A and D748Q in strand β1 and W768V and C773S in strand β3), and the chimeric construct resulting from the combination of ADAR1-dsRBD3 N- and C-terminal fragments flanking the Xlrbpa-dsRBD2 domain. Residue numbering and secondary structure elements of ADAR1-dsRBD3 are shown above the alignment. Identical residues between ADAR1-dsRBD3 and Xlrbpa-dsRBD2 are highlighted *in green*. The four mutations introduced in ADAR1-dsRBD3 in order to disrupt the dimer interface (i.e. V747A, D748Q, W768V and C773S) are shown *in yellow*. The three mutations introduced in Xlrbpa-dsRBD2 in order to adapt the surface of Xlrbpa-dsRBD2 for enabling the tight interaction of helix αN in between helices α1 and α2 (i.e. S116G, T176R and K179I) are shown *in blue*.

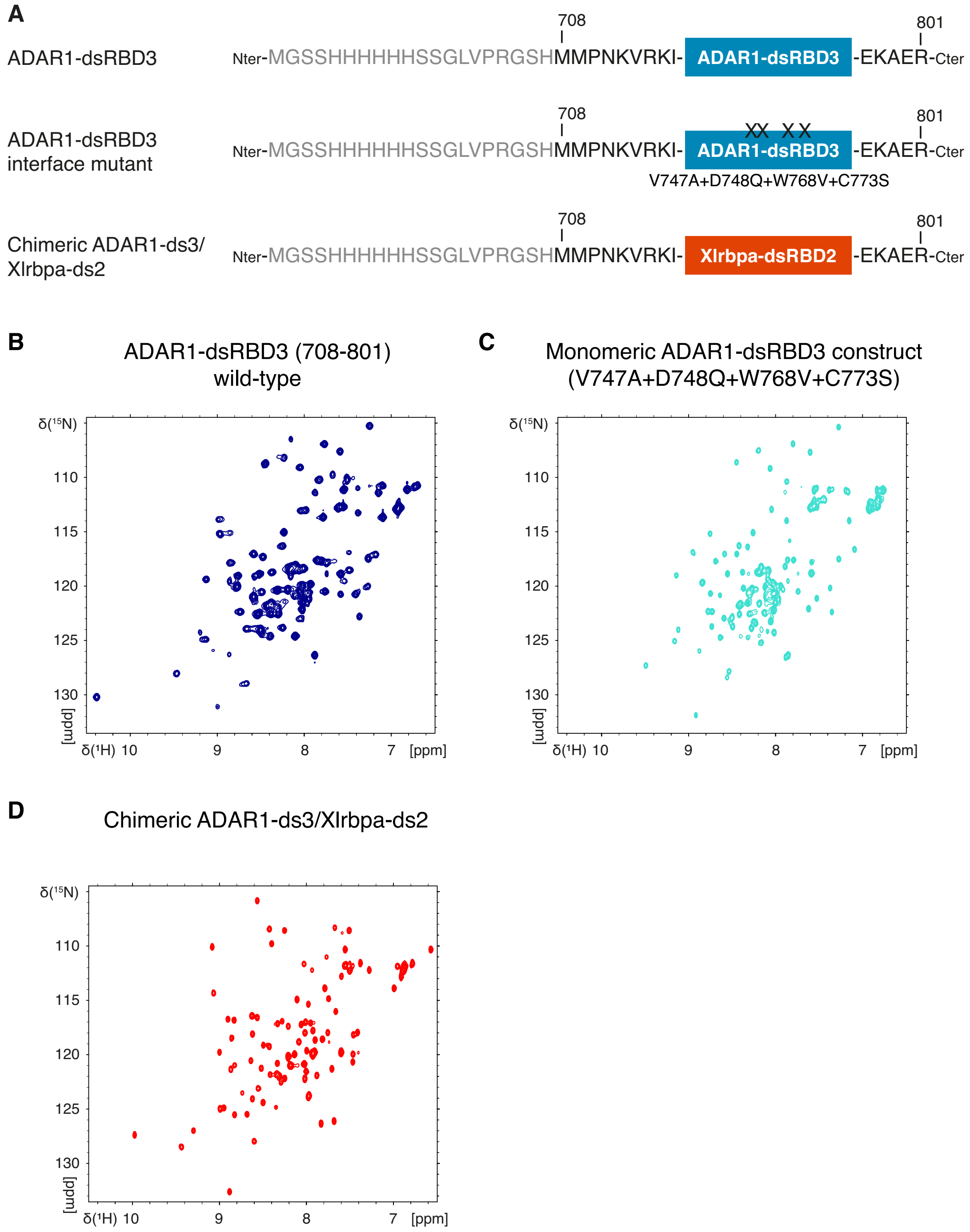

**Supplementary Figure S5: Monomeric ADAR1-dsRBD3 mutant and chimeric ADAR1-dsRBD3/Xlrbpa-dsRBD2 are well-folded domains.**

(**A**) Schematic representation of ADAR1-dsRBD3 constructs: ADAR1-dsRBD3 (residues 708-801), ADAR1-dsRBD3 interface mutant (residues 708-801; V747A+D748Q+W768V+C773S), and chimeric ADAR1-ds3/Xlrbpa-ds2 (see Supplementary Figure S4). Constructs used here retained their purification tags. Residues of the N-terminal $His_6$-tag are shown in grey. (**B**) ($^{1}H,^{15}N$)-HSQC spectra from ADAR1-dsRBD3 wild-type (*in deep blue*). (**C**) ($^{1}H,^{15}N$)-HSQC spectra from monomeric ADAR1-dsRBD3 (mutant V747A+D748Q+W768V+C773S) (*in light blue*). (**D**) ($^{1}H,^{15}N$)-HSQC spectra from chimeric ADAR1-dsRBD3/Xlrbpa-dsRBD2 (*in red*). The dispersion of amide signals in all constructs (panel B-D) is the sign of well-folded domains.

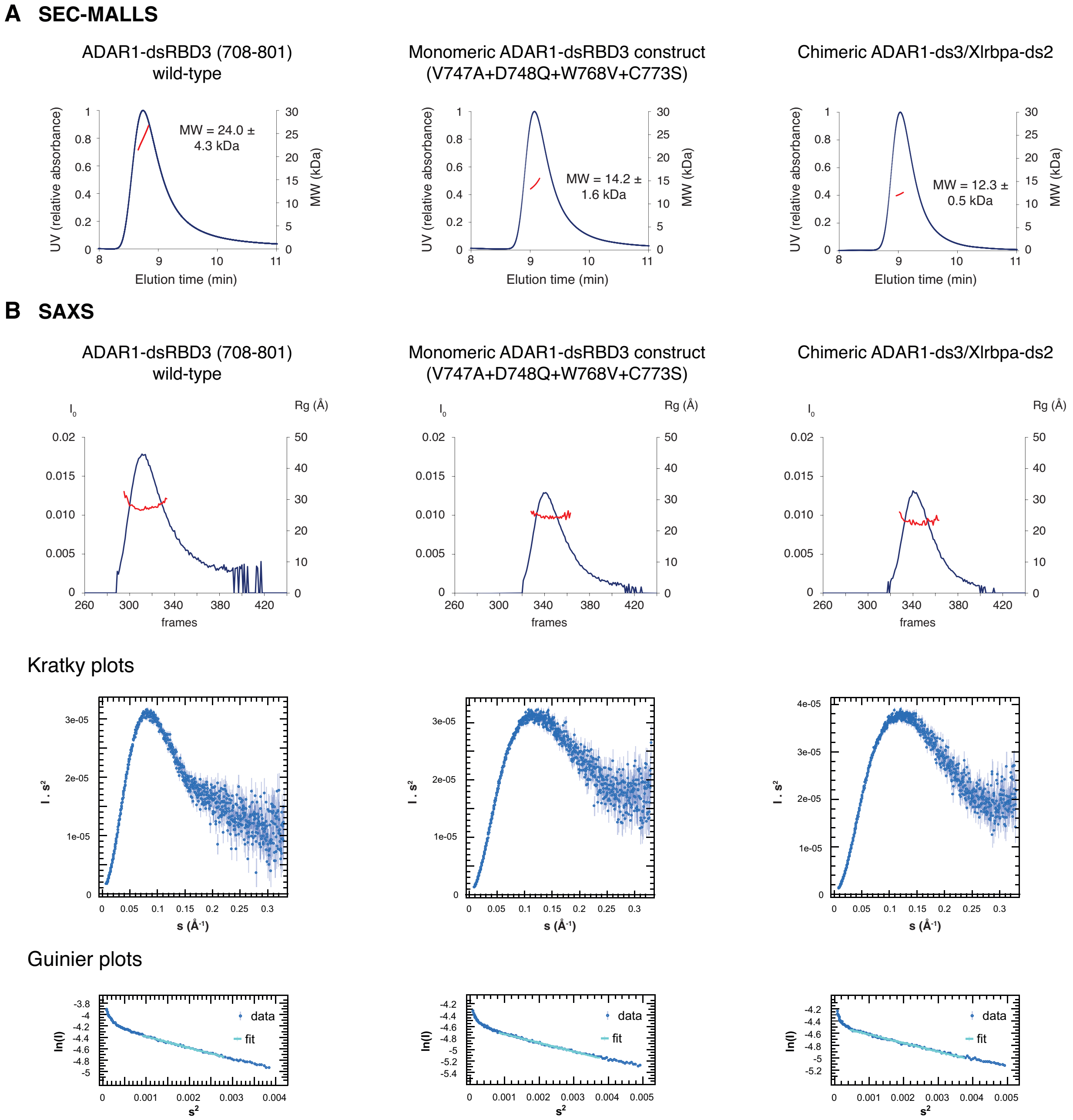

**Supplementary Figure S6: Characterization of monomeric ADAR1-dsRBD3 and chimeric ADAR1-dsRBD3/Xlrbpa-dsRBD2 constructs by SEC-MALLS and SAXS.**

SEC-MALLS and SAXS data are presented for the following ADAR1-dsRBD3 constructs: ADAR1-dsRBD3 wild-type (residues 708-801), ADAR1-dsRBD3 interface mutant (residues 708-801; V747A+D748Q+W768V+C773S), and chimeric ADAR1-ds3/Xlrbpa-ds2. Constructs used here retained their purification tags (see Supplementary Figures S4 and S5). (**A**) SEC-MALLS analysis of the constructs on a BioSEC-3 column. The UV relative absorbance and the estimated molecular weight (MW) obtained around the elution peak of HPLC are displayed in blue and in red, respectively. Molecular weights values are directly reported on the figure for each construct. (**B**) $I_0$ and Rg values obtained around the elution peak of HPLC are displayed in the upper panel. Kratky plots are shown in the middle panel (Kratky plots). Regions used for the Guinier approximation are shown in the lower panel (Guinier plots). Derived parameters are reported on Table 2.

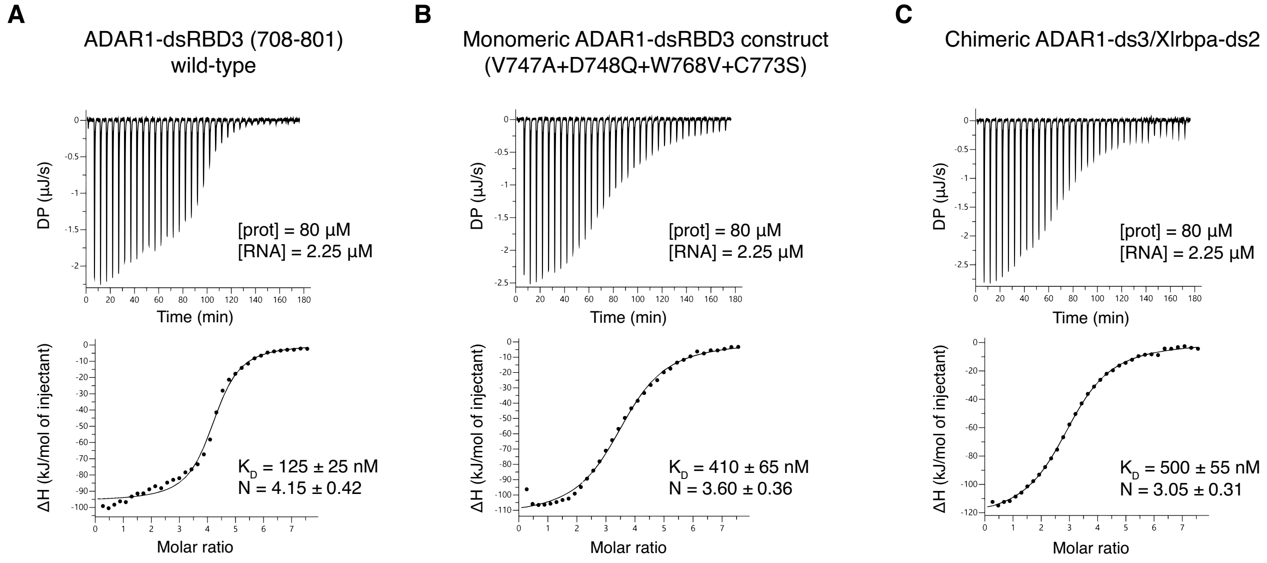

**Supplementary Figure S7: Monomeric ADAR1-dsRBD3 remains competent for RNA-binding.**

Isothermal titration calorimetry monitoring the interaction between ADAR1-dsRBD3 wild-type (**A**), monomeric ADAR1-dsRBD3 construct (**B**), and chimeric ADAR1-dsRBD3/Xlrbpa-dsRBD2 construct (**C**) and a 24 bp dsRNA duplex. $K_D$: dissociation constant in nM. N: number of sites. Values are reported as means ± standard error (S.E.). The uncertainties on the fitted parameters were estimated from the data spread and from the uncertainty of the protein concentration determination (10%).

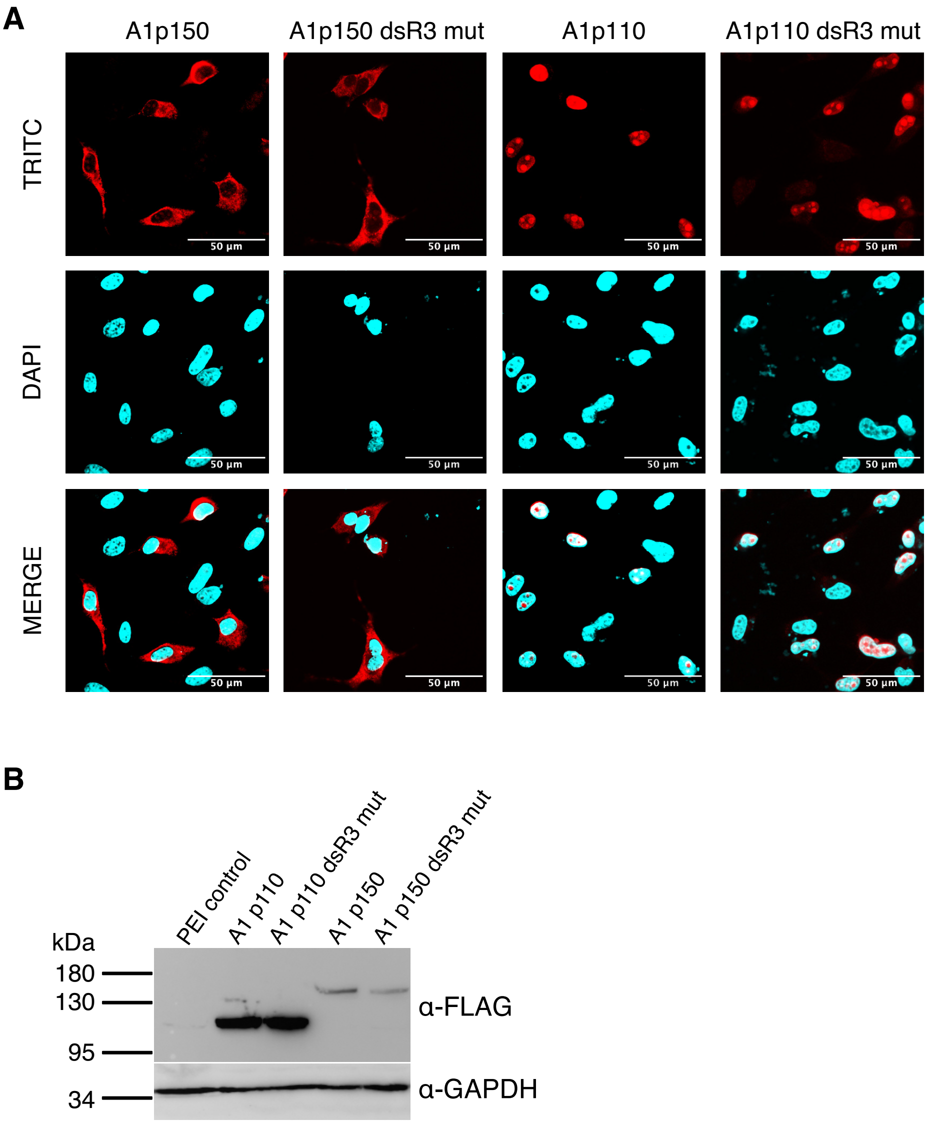

**Supplementary Figure S8: ADAR1-dsRBD3 interface mutations do not affect cellular localization of ADAR1 isoforms p110 and p150.**

(**A**) Confocal microscopy images confirm the cellular localization of ADAR1 variants. Confocal sections of transfected constructs are visualized under TRITC channel. DAPI is used for nuclear staining (scale bar: 50 µm). (**B**) Western blots confirm the similar level of expression of ADAR1 p110 (A1 p110) with respect to its dsRBD3 dimerization mutant (A1 p110 dsR3 mut) and ADAR1 p150 (A1 p150) with respect to its dsRBD3 dimerization mutant (A1 p150 dsR3 mut). Expression of proteins detected by anti-FLAG antibodies. GAPDH serves as a loading control and PEI control indicate transfected cells without expression plasmids.
